# The language network ages well: Preserved topography, lateralization, selectivity, and within-network functional connectivity in older brains

**DOI:** 10.1101/2024.10.23.619954

**Authors:** Anne Billot, Niharika Jhingan, Maria Varkanitsa, Agata Wolna, Cory Shain, Idan Blank, Rachel Ryskin, Swathi Kiran, Evelina Fedorenko

## Abstract

Healthy aging is associated with structural and functional brain changes. However, cognitive abilities vary in how they change with age: whereas executive functions, like working memory, show age-related decline, aspects of linguistic processing remain relatively preserved. The heterogeneity of the cognitive-behavioral landscape in aging predicts differences among brain networks in whether and how they should change with age. To evaluate this prediction, we used individual-subject fMRI analyses (’precision fMRI’) to examine the language-selective network and—for control purposes—the Multiple Demand (MD) network, which supports executive functions, in older adults (n=64) relative to young controls (n=483). In line with past claims, relative to young adults, the MD network of older adults shows weaker, less spatially extensive, and more topographically variable activations during an executive function task and reduced within-network functional connectivity. However, in stark contrast to the MD network, we find remarkable preservation of the language network in older adults. Their language network responds during language comprehension as strongly and selectively as in younger adults, and shows a similar degree of left-hemispheric lateralization and within-network functional connectivity. Our findings suggest that the language network remains young-like—at least on standard measures of function and connectivity—and align with behavioral preservation of language comprehension in healthy aging.

## Introduction

Healthy aging is associated with slower information processing and a reduction in working memory and attentional resources ^1–4^. These behavioral changes in executive abilities have been argued to result from changes in the structure and function of different large-scale brain networks. Some have reported age-related changes in the brain areas that support executive functions (e.g., ^5–13^). Others have argued for more widespread effects on multiple networks. For example, one popular claim has been that brain networks become generally less clearly segregated with age^14–16,16–25^. This reduced segregation can manifest as less clearly distinguishable functional profiles—de-differentiation^26^—due to a combination of weaker responses in functionally specialized areas and compensatory recruitment of brain regions not typically engaged in those functions in younger individuals (often, the frontal executive regions^5,26^). Reduced segregation can also manifest as weaker dissociations in functional connectivity, with reduced within-network connectivity and increased between-network connectivity^14–25^.

In contrast to the well-documented changes in executive abilities ^1–4^, language is often described as one of the better-preserved functions in older adults. In spite of occasional word-finding difficulties (e.g., ^27,28^; for a review, see ^29^) and changes in the organization of semantic memory ^30–35^, vocabulary keeps growing with age, the ability to extract meaning from texts continues to improve, and the ability to predict upcoming words remains relatively intact^2,28,36–43^ (for reviews, see ^26,44–48^). However, whether and how the neural infrastructure for language processing changes with age remains debated. For example, some researchers have argued that the general topography of the language network—a set of temporal and frontal brain areas that respond selectively during language processing^49^—is similar between younger and older adults^45,50^, but others have argued for more bilateral activation with age^51–53^. Similarly, the magnitude of response in some language regions has been argued to increase with age^51,54,55^, to decrease with age^37^, or to show no change^37,50,51,56^. And for within-network connectivity, some have reported a general reduction with age^57,58^, others—an increase or a decrease depending on the specific pair of brain regions^28^, yet others found no change^50,59^.

These inconsistencies could be due to several factors. First, past studies have used an array of language tasks—from object naming^37,52,60^, to auditory sentence comprehension ^50,54^, to monitoring of a target word^51^, to passive narrative reading^56^—making across-study comparisons challenging. Moreover, many commonly used paradigms conflate language processing and general task demands^61^. Because such paradigms recruit both the language-selective network and the domain-general Multiple Demand (MD) network, which supports executive functions^62,63^, the observed effects are difficult to attribute to a particular network. This issue is compounded by the reliance of most past fMRI studies on group analyses, where activation maps or functional connectivity maps are averaged voxel-wise across participants^51,57,58,60,64–67^. Because of substantial inter-individual variability in the precise locations of functional areas^68–72^, including language areas^73–76^ and MD areas^63,69,77–80^, and the proximity of the two networks to each other in the association cortex^77,79^, this group-averaging approach blurs network boundaries leading to interpretive challenges^61,81,82^. This blurring of functional boundaries, inherent in the group-analytic approach, is the reason that the field of cognitive neuroscience has been increasingly moving toward ‘precision imaging’^61,77,83–85^, where the relevant functional areas are identified within individuals, either using established ‘localizer’ paradigms or bottom-up clustering of voxel timecourses based on functional connectivity data (^69,73,77,80^; for general discussions of the advantages of precision fMRI, see ^61,83,84^).

To resolve some of the controversies about whether or how the language network changes with age, we here use a precision imaging approach with a ‘localizer’ paradigm that has been established in prior studies to reliably identify the language network, which selectively supports comprehension and production (for a review see ^49^). For comparison, we also examine the domain-general Multiple Demand (MD) network^86^, which supports executive abilities, such as attention, working memory, and cognitive control^62,86–88^—functions that are widely agreed to exhibit age-related decline^89–97^. Examining these two cognitive networks in parallel enables a direct test of whether age-related brain changes are uniform across networks or instead exhibit network-specific profiles. In addition, this approach allows us to evaluate the claim about network de-differentiation introduced above^26^. In line with increasing emphasis in the field on robustness and replicability^98–100^, we i) examine a relatively large cohort of older adults (OA, n=64), along with a large cohort of young adults (YA, n=483), ii) include several commonly used measures of brain function, and iii) use several statistical analytic approaches. To foreshadow our results, we find that a) for the language network, the topography, including the extent of activation and degree of lateralization, as well as the strength of within-network connectivity are similar between older and younger adults, and only the magnitude of response to language slightly increases with age, and b) for the MD network—which remains robustly segregated from the language network in older adults—the topography is more variable across individuals and the magnitude and extent of activation as well as the within-network connectivity decrease substantially with age.

## Results

Our participants consisted of 64 older adults (OA) (average age: 59.7; range 41-80; 36 females; 59 right-handed, 3 left-handed, one ambidextrous and one with missing handedness data) and 483 younger adults (YA) (average age: 23.8; range 17-39; 278 females; 431 right-handed, 28 left-handed, 11 ambidextrous and 13 with missing handedness data). The OA cohort includes data from two independent cohorts (**Figure 1A**; <u>Methods</u>) that were combined to increase statistical power (but see cohort-specific results in **Figures 2-3** and in **Supplementary Tables 3** and **6**). General cognitive functioning was assessed in one of the OA cohorts (OA2) using the MMSE-2^101^, and all participants scored within the normal range (i.e., within 1 standard deviation from the mean). Each participant completed two extensively validated localizer tasks (**Figure 1B-C**), which reliably and selectively engage the target networks of interest—the language and the MD networks—in spite of their close proximity within the left frontal lobe^77,102–105^ (see **Figure 2A** for sample individual activation maps). A subset of the participants additionally completed one or two so-called ‘naturalistic-cognition’ paradigms: a resting state scan and a story listening paradigm. These data were used in the analyses of functional connectivity, and the story listening data were used to test for sensitivity to linguistic complexity in the language network’s responses.

**Figure 1.**
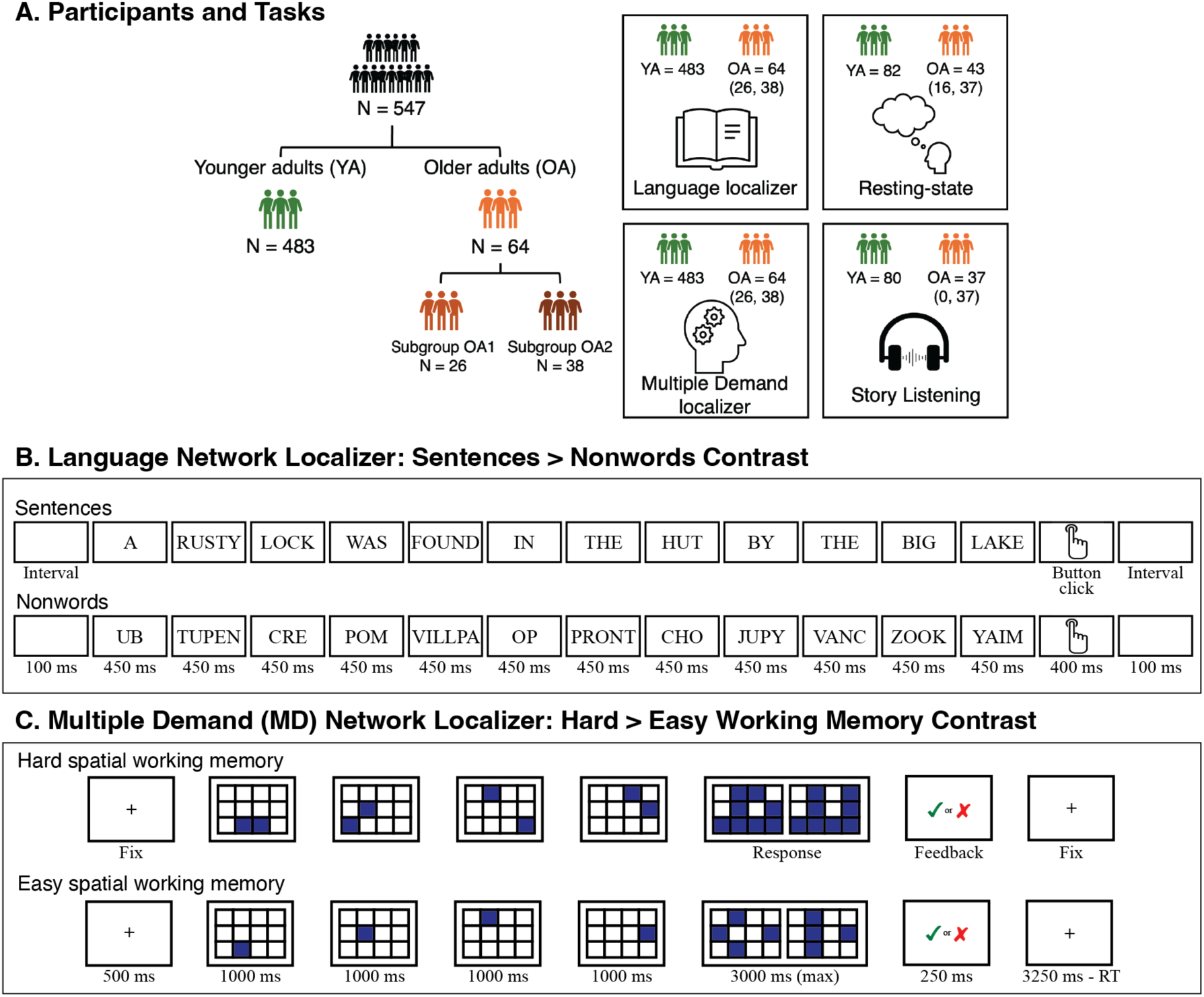
Participants and task paradigms. A) The dataset includes a large cohort of young adults (YA), and two cohorts of older adults (OA), which are combined in all the main analyses (<u>Methods</u>). On the right, the numbers in parentheses indicate the number of participants from each OA cohort (OA1, OA2) who completed each task. B) In the language localizer ^73^, participants were asked to attentively read sentences (critical condition) and lists of pronounceable nonwords (control condition) in a blocked design, one word or nonword at a time, and to press a button at the end of each sentence/nonword-list. C) In the MD localizer (a spatial working memory task; ^63,126^), participants were asked to keep track of eight (hard, critical condition) or four (easy, control condition) spatial locations (presented two at a time, or one at a time, respectively) in a 3 × 4 grid, in a blocked design. At the end of each trial (in both conditions), participants were asked to perform a two-alternative forced-choice task to indicate the set of locations they just saw. (See <u>Methods</u> for additional details of the paradigms.)

**Figure 2.**
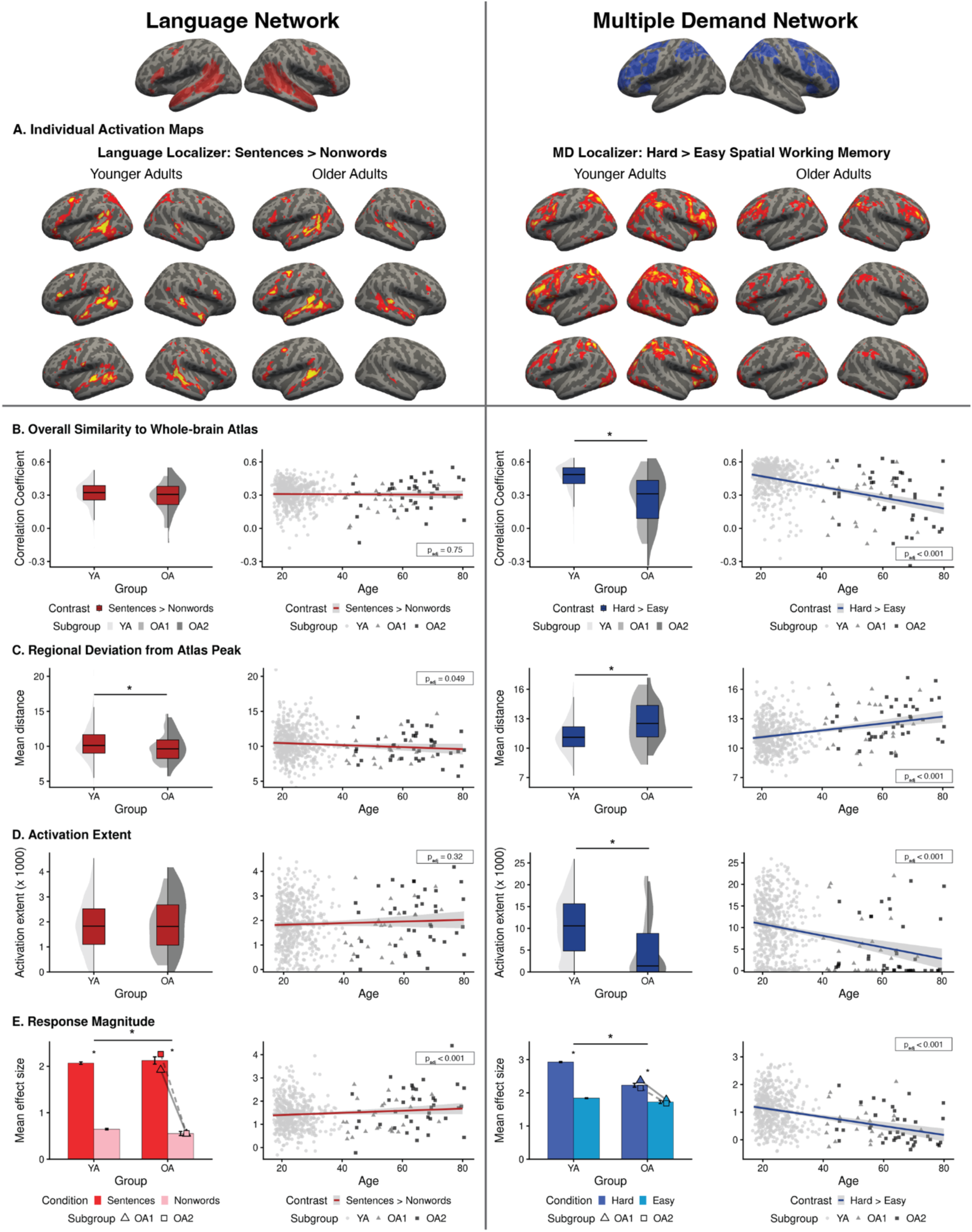
Dissociable effects of aging on the language network (left) and the Multiple Demand (MD) network (right). **A)** Sample individual activation maps for the two localizer contrasts. The maps are thresholded at the whole-brain level of *p*<0.01 (uncorrected) and are included solely for illustrative purposes; all statistical analyses were performed on neural measures extracted from these maps as described in <u>Methods</u>. The full set of individual maps is available on OSF (https://osf.io/2q65t/?view_only=ab1833db12c64eb0a7cc61c5795d35cd). Neural responses in young adults (YA) and older adults (OA) are compared for four neural measures. Across these measures, the language network shows preserved function with age (if anything, activations appear to be slightly less variable and stronger in OA, compared to YA), whereas the MD network shows consistent decline. **B)** Global topographic similarity of the bilateral language and MD networks measured as the spatial correlation between an individual’s activation map and the corresponding probabilistic network atlas. **C)** Regional deviation of activation peak locations in the LH language and bilateral MD networks measured as the mean Euclidean distance (in mm) between an individual’s activation peak and a normative probabilistic atlas centroid within functional parcels (or “search spaces”). **D)** Extent of activation measured as the number of significant voxels for the relevant contrast at the FDR-corrected *p*<0.05 threshold within each network’s parcels for the LH language and bilateral MD networks. **E)** Mean BOLD response magnitude during the language task in the LH language network and during the spatial working memory task in the bilateral MD network. All fROIs are defined within individual participants, and responses are estimated using across-runs cross-validation (see <u>Methods</u>). The OA group is divided into two cohorts (OA1 – triangles, and OA2 - squares). Significant positive task contrast effects (*p_adj_*<0.05) are marked with an asterisk between bars. Significant group differences (*p_adj_*<0.05) are marked with an asterisk above a line. In all scatter plots, individual dots represent participants (YA: light grey circles; OA1: grey triangles; OA2: dark grey squares), the solid line is the linear regression fit, and the shaded area is the 95% confidence interval. Error bars on bar plots represent SEM.

**Figure 3.**
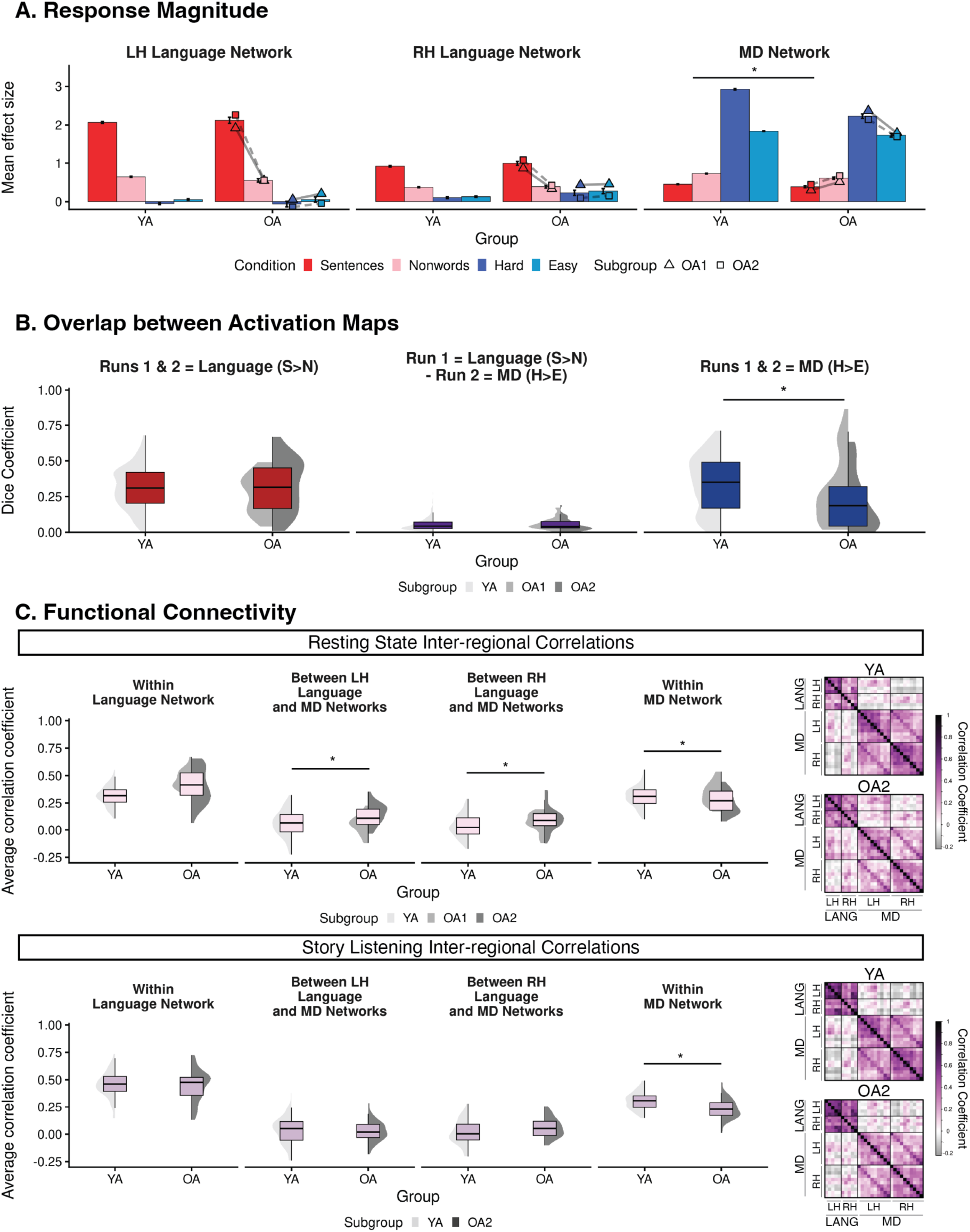
Preserved functional dissociation between the language network and the Multiple Demand (MD) network with age. The older adult (OA) group is divided into two cohorts (OA1 – triangles, and OA2 - squares) and shows no significant difference from the younger adults (YA) group in responses to the non-preferred domain, activation overlap between networks, or between-network connectivity during language processing. **A.** Mean BOLD response to the conditions of the language task (sentence reading: red bars; nonword reading: pink bars) and the spatial working memory task (hard condition: dark blue bars; easy condition: light blue bars) in the left hemisphere (LH) and right hemisphere (RH) language networks and in the bilateral MD network (all fROIs are defined within individual participants, and responses are estimated using across-runs cross-validation; see <u>Methods</u>). Significant positive task contrast effects (*p_adj_* < 0.05) are marked with an asterisk between bars. Significant differences between groups (*p_adj_*<0.05) in the task contrast effects for the non-preferred domain (i.e., the hard vs. easy spatial working memory effect in the language fROIs, and the sentences vs. nonwords effect in the MD fROIs) are marked with an asterisk above a line (see Figure 2 for group differences in response to the preferred domain). **B.** Spatial overlap of significantly activated voxels across runs within a localizer task (sentences > nonwords: left, hard > easy: right) and between the two tasks (middle), as measured with the Dice coefficient. Whole-brain activation maps were thresholded at FDR-corrected *p*<0.05. **C.** <u>Box plots</u>: Average inter-regional timeseries correlations among the regions of the language network (left), among the regions of the MD network (right), and between the language and MD networks (two middle panels) during a resting-state paradigm (top row) and story listening (bottom row). <u>Correlation Matrices</u>: Inter-regional functional correlation matrices for pairs of regions within each network and between the two networks during a resting-state paradigm (top row) and story listening (bottom row) in the YA and OA2 groups. LH: left-hemisphere fROIs, RH: right-hemisphere fROIs; LANG: language, MD: Multiple Demand. Significant group differences (*p_adj_*<0.05) are marked with an asterisk above a line. Error bars on bar plots represent SEM.

Before presenting the critical results, we describe the two brain networks of interest in more detail. The language network, as defined here, is a set of left-lateralized frontal and temporal brain regions that jointly support linguistic processing, including lexical retrieval, syntactic structure building, and semantic composition (^73,106–109^; see ^49^ for a review). Although this network necessarily interacts with other brain systems during real-life language use, its regions are strongly selective for language over diverse non-linguistic inputs and tasks (^77,110–115^; see ^49,116^ for reviews). The Multiple Demand (MD) network is a set of bilateral frontal and parietal brain regions that jointly support executive functions, including attention, working memory, and cognitive control^62,86,87^. The MD network is engaged by diverse cognitively demanding tasks, across stimulus and task types^63,117–120^. It is important to note that although we use particular localizers to define our networks of interest, each of these paradigms has been shown to generalize across many variants that use alternative materials and tasks ^63,119,121–123^. Furthermore, both of these networks can also be defined in individual participants based on patterns of functional connectivity alone^69,77,124,125^ (for earlier relevant evidence, see ^103^), and the networks thus identified correspond closely with the activations elicited by validated localizers, like those used here (see **Supplementary Figure 2a** for evidence from the current dataset).

In the analyses, we focus on the neural measures that have been most commonly examined in past studies. In particular, for both networks, we first consider the typicality of the activation patterns by comparing individual maps to probabilistic activation atlases from large numbers of participants scanned on the same paradigms in prior studies. We additionally examine changes in the extent of activation and, for the language network only, the degree of lateralization. Lastly, we investigate the magnitude of neural response for the relevant contrasts and the strength of inter-regional functional correlations during the naturalistic paradigms. To ensure robustness to analytic choices, we report both categorical effects of age group, and continuous effects of age across the full set of participants.

### 1. The language network is well-preserved in aging

The language network of older adults was remarkably similar to that of younger adults across diverse neural measures, including measures of activation topography, response magnitude, and functional connectivity.

First, we examined the topography of the network, to test whether it may become less typical or more variable with age. To evaluate the overall typicality of the activation landscape, we compared whole-brain individual language activation maps (see **Figure 2A** for sample maps) to the probabilistic language network atlas created from 806 activation maps (Lipkin et al., 2022). We found no difference between the age groups, nor as a function of continuous age (**Figure 2B left panel; Supplementary Table 3.1.1**). To test the degree of inter-individual variability in the locations of regional activation peaks, we computed Euclidean distances between each individual’s peak and the peak in the probabilistic atlas for the 5 left-hemisphere (LH) language regions (**Figure 2 top left panel**). The variability in the peak locations was, on average, slightly lower in OA compared to YA (**Figure 2C left panel; Supplementary Table 3.1.2**; OA>YA group effect: *β*=-0.585, t=-2.03, *p_adj_*=0.043; continuous age effect: *β*=-0.014, t=-1.97, *p_adj_*=0.049). Finally, the language network of OA remains strongly lateralized to the left hemisphere, with no significant difference relative to the YA group or as a function of continuous age (**Supplementary Table 3.1.3**).

Next, we examined the extent and magnitude of activation for the language localizer contrast in the core LH areas. We found no difference between the YA and OA groups in the extent of activation, nor as a function of continuous age (**Figure 2D left panel; Supplementary Table 3.1.4;** see **Supplementary Figure 4A** for the effects broken down by fROI and **Supplementary Table 6.2.1** for similar results at the whole-brain level). We did, however, observe a slightly higher response to the language contrast (i.e., sentences > nonwords) in the OA relative to the YA group (**Figure 2E left panel** and **Supplementary Table 3.1.5;** OA>YA group effect: *β*=0.082, t=2.78, *p_adj_*=0.012; continuous age effect: *β*=0.003, t=4.29, *p_adj_*=0.002). The difference is small (the average sentences>nonwords effect in the YA is 90.5% of the effect size in the OA), and the effect appears to be mostly restricted to the left posterior temporal region (**Supplementary Figure 4b**, OA>YA group effect False Discovery Rate (FDR)-corrected *p*<0.020; continuous age effect: FDR-corrected *p*=0.001). We come back to this difference in the <u>Discussion</u>. In the RH homotopic network, the YA and OA groups did not show any significant differences in the magnitude of response to the sentences>nonwords contrast (**Supplementary Figure 5**; this figure also shows the results of exploratory analyses of brain areas in the extended language network ^127^).

Finally, the degree of functional connectivity among the regions of the LH language network, as measured by Pearson’s correlations between BOLD signal timeseries across pairs of language fROIs, was similar between the YA and OA groups during both rest and story comprehension (**Figure 3C; Supplementary Table 3.1.6**). The results were also similar for the RH homotopic network (**Supplementary Table 3.1.6**) and for pairs of fROIs between hemispheres (**Supplementary Tables 6.3.1** and **6.3.2**).

The language localizer paradigm in the current study used passive comprehension. Because much evidence implicates language-selective brain areas in computations related to word retrieval and combinatorial linguistic processing (e.g., ^108^), strong responses in these areas suggest that participants were engaging these computations. However, to more directly test such engagement—and to evaluate our participants’ comprehension ability—we leveraged the data from the naturalistic story paradigm, which was administered to the OA2 cohort. First, we tested whether neural responses to language in the older adults show canonical signatures of language processing—sensitivity to word frequency and surprisal, with lower-frequency and more contextually unexpected words being more costly to process (behavioral evidence: ^128,129^, neural evidence: ^130–132^). To do so, we modeled responses to each word during the naturalistic story comprehension task as a function of frequency and surprisal (estimated using GPT2-xl;^133^), with controls for word rate (a regressor at the onset of each word) and end-of-sentence markers. This model revealed reliable effects of frequency and surprisal in the language areas (*p*s<0.001), mirroring the findings previously reported for young adults ^132,134^. Next, we evaluated comprehension ability by analyzing responses to a set of questions administered after the story to the OA2 cohort (<u>Methods</u>). The accuracies were high, with almost all participants answering all questions correctly (average = 97%, st. dev. = 10.3%). These results suggest that the language areas of older adults are processing linguistic input in a similar way to those of younger individuals, showing strong sensitivity to linguistic complexity, and that comprehension ability is strong in our OA participants, in line with past evidence of intact comprehension ability in healthy aging ^2,36–41,135^.

### 2. The Multiple Demand network shows pronounced age-related changes

In contrast to the preservation of the language network in older adults, we expected to find age-related changes in the Multiple Demand (MD) network given that this network supports executive functions, which decline with age ^2,136–138^. Indeed, in line with prior studies ^5,8–13,18,19,37,139,140^, we observed age effects across all neural measures examined.

First, we compared individual MD activation maps to the probabilistic MD network atlas created from 691 activation maps with respect to overall topography and activation peak locations. We found that the topography of the MD network was more variable in older individuals as evidenced by a) lower correlations between whole-brain activation maps and the MD atlas (**Figure 2B right panel; Supplementary Table 3.2.1**; OA>YA group effect: *β*=-0.18, t=-9.16, *p_adj_*<0.001; continuous age effect: *β*=-0.005, t=-9.97, *p_adj_*<0.001), and b) larger Euclidean distances between individual peaks and the peak in the probabilistic atlas for the 20 MD regions (10 in each hemisphere) (**Figure 2C right panel; Supplementary Table 3.2.2**; OA>YA group effect: *β*=1.416, t=6.35, *p_adj_*<0.001; continuous age effect: *β*=0.035, t=6.23, *p_adj_*<0.001).

Next, we examined the extent and magnitude of activation for the localizer contrast in the MD areas. The older adults showed less extensive activation during the spatial working memory task compared to younger adults. This effect held both when restricting the analysis to the MD parcels that cover broad areas of the typical MD network response (**Figure 2D right panel; Supplementary Table 3.2.3**; OA>YA group effect: *β*=-9.09, t=-7.22, *p_adj_*<0.001; continuous age effect: *β*=-0.23, t=-7.34, *p_adj_*<0.001) and when considering whole-brain activations (**Supplementary Table 6.2.1.**; OA>YA group effect: *β*=-18.07, t=-2.37, *p_adj_*=0.004; continuous age effect: *β*=-0.50, t=-3.13, *p_adj_*=0.004).

Mirroring the activation extent effects, the OA group showed significantly lower responses to the spatial working memory task contrast (i.e., hard > easy) in the MD fROIs. This effect held both at the network level (**Figure 2D right panel; Supplementary Table 3.2.4**; OA>YA group effect: *β*=-0.33, t=-26.52, *p_adj_*<0.001; continuous age effect: *β*=-0.009, t=-15.16, *p_adj_*<0.001) and in each MD fROI individually (**Supplementary Figure 4b**; OA>YA group and continuous age effects: FDR-corrected *ps*<0.001). Post-hoc analyses revealed that this reduction in the size of the hard > easy effect in the OA group was driven by a lower response during the hard condition (OA>YA group effect: *β*=-0.43, t=-5.70, *p*<0.001; continuous age effect: *β*=-0.01, t=-5.96, *p*<0.001); in contrast, the groups did not differ in their response during the easy condition. It therefore appears that with age, the MD network’s ability to increase activity in response to higher cognitive demands decreases. These neural changes were accompanied by a decline in behavioral performance, with older adults showing lower accuracies for both the hard and easy conditions compared to the YA group (**Supplementary Figure 1b**; *p*s<0.001). However, even after accounting for individual differences in task performance, the analysis with age as a continuous variable (performed on the YA and OA1 cohorts, where an identical version of the task was used) revealed a significant effect of age on the hard>easy contrast magnitude (*β*=-0.006, t=-2.03, *p*=0.043, **Supplementary Table 3.2.5**). These brain-behavior analyses also revealed that for both YA and OA1, better task performance is associated with lower responses during the easy trials and higher responses during the hard trials (see ^126^ for concordant evidence from younger adults).

Finally, functional connectivity among the regions of the MD network, as measured by Pearson’s correlations between BOLD signal timeseries across pairs of MD fROIs, was weaker in the OA group (**Figure 3C**) during both rest (OA>YA group effect: *β*=-0.08, t=-4.38, *p_adj_*<0.001; continuous age effect: *β=*-0.003, t=-5.2, *p_adj_*<0.001) and story comprehension (OA>YA group effect: *β*=-0.06, t=-3.35, *p_adj_*=0.017; continuous age effect: *β*=0.002, t=-3.75, *p_adj_*<0.001). These results also held when considering fROI pairs within each hemisphere separately (**Supplementary Table 3.2.6**: all *p_adj_* < 0.007).

### 3. The language and the MD networks remain robustly segregated in older adults

Finally, to evaluate prior claims about functional networks becoming less clearly differentiated with age^14–25^, we examined the relationship between the language and the MD networks across several measures.

First, the functional profiles of the language and MD regions remain clearly distinct with age. In both younger and older adults, the language network shows an overall low response during the spatial working memory task and does not show a significant hard > easy effect either at the network level (**Figure 3A left and middle panels**), or in any individual language region (**Supplementary Figure 4b** top panel). The MD fROIs show an overall low response during language processing and show the opposite effect compared to the language network: a stronger response to nonword-lists than meaningful sentences. No significant differences were found between YA and OA in the responses to the spatial working memory task in either hemisphere of the language network (**Figure 3A left and middle panels; Supplementary Table 3.3.1**). In the MD network, on average, OA showed numerically lower responses to both conditions of the language task compared to YA, but the differences were small (sentences: mean YA = 0.45, mean OA = 0.37; nonwords: mean YA = 0.73, mean OA = 0.59). The nonwords>sentences effect became slightly smaller in OA compared to YA (**Figure 3A right panel**, YA > OA group effect: *β*=0.039, t=2.83, *p_adj_*=0.028) but did not come out as significant as a function of continuous age, after correction for multiple comparisons (*β*=0.001, t=2.42, *p*=0.015, *p_adj_*=0.076) or when breaking the MD network by hemisphere (**Supplementary Table 3.3.1**).

Second, the OA group did not differ from the YA group in the degree of spatial overlap between the thresholded individual activation maps for the language and the MD localizer contrasts. The overlap, as measured by the Dice coefficient^141^, was minimal for both groups (**Figure 3B middle panel; Supplementary Table 3.3.2**) and significantly lower than the overlap across runs within either network for their respective localizer task contrast (**Figure 3B left and right panels**, all *p_adj_*<0.001).

Finally, similar to the YA group, the OA group showed a clear dissociation between the language and the MD networks in the functional connectivity patterns, with higher within-network than between-network connectivity (**Figure 3C**; all *p_adj_*<0.001). We did observe a small but significant increase in the between-network connectivity during rest in the OA group (OA>YA group effect: *β*=0.062, t=4.23, *p_adj_*<0.001; continuous age effect: *β*=0.002, t=3.81, *p_adj_*<0.001, **Supplementary Table 3.3.3**). However, this effect was not present during the story listening task, ruling out an explanation of a compensatory mechanism for language processing, and raising questions about the robustness of this age effect across cognitive states.

## Discussion

We used individual-subject fMRI analyses (’precision fMRI’^82^) and extensively validated ‘localizer’ paradigms to examine age-related changes in the language-selective network and, for comparison, in the domain-general Multiple Demand (MD) network. Our findings reveal a striking dissociation in the impact of aging on these two cognitive networks. The language network shows remarkable preservation with age, reflected in similar topography—including typicality (similarity to the prototypical activation pattern) and similar amount of inter-individual variability in regional peak locations—and lateralization, similarly strong responses to a language comprehension task, and similarly strong within-network functional connectivity. These results align with relative preservation of linguistic comprehension abilities in aging^2,26,36–38,44–47^. In contrast, for all neural measures examined, the MD network showed age-related decline: the activation topography was less typical and more variable, the responses during a working memory task were reduced, and the within-network functional connectivity was lower in older adults, in line with age-related decay in executive abilities^89–97,142^, which the MD network supports. In the remainder of the <u>Discussion</u>, we contextualize these findings with respect to the prior literature on cognitive aging and discuss their implications.

### Preservation of the language network with age

As outlined in the Introduction, language processing—which includes the retrieval of words from memory, syntactic structure building, and semantic composition during comprehension and production—is one of the better-preserved cognitive functions in older age^26,36–38,44–47^. In line with this behavioral preservation, in a large sample of participants, we found that the language-selective network, which supports our ability to interpret and generate linguistic messages across modalities^49,143–147^ is preserved with age. These results held regardless of whether age was treated as a categorical or continuous predictor.

The only small but significant difference we observed between age groups within the language network was a slightly higher response to the language localizer contrast (i.e., reading sentences > nonwords) in older adults. The level of response in the language network has been shown to scale with how competent the individual is as a language user. For example, responses increase across the developmental trajectory, between age 4 and late adolescence, at which point they asymptote^148–150^. Similarly, responses are stronger in adult non-native speakers who are more proficient in the language compared to those who are less proficient^151^. These effects are presumably due to the fact that children and adults with limited proficiency may not be able to engage linguistic computations to the full degree because they may be unfamiliar with certain words or constructions, leading to the inability to piece together the complete meaning. Older native speakers, such as the participants in the current study, are of course highly proficient language users: the ability to interpret and generate linguistic messages does not appear to decay with age, except for occasional word-finding difficulties^2,36–43^; for reviews, see ^26,44–48^). In fact, vocabulary size continues to increase with age ^32,152,153^, which may explain the slightly higher responses to language that we observed in older adults. In addition, when listening to a naturalistic story—edited to include rare words and constructions^154,155^—the language areas of older adults exhibit canonical signatures of language processing: increased response to infrequent and contextually surprising words (e.g., ^130,132^). This result argues against qualitative age-related changes in how linguistic inputs are processed in this network.

The preservation of the language network is consistent with the brain maintenance hypothesis^156,157^, whereby healthy aging, as measured by the stability of cognitive performance in some domain over the adult life course, is due to the preservation of the structure and function of the relevant neural substrate^156–158^. Prior evidence for this hypothesis has primarily come from individual-differences investigations in the domains of long-term episodic memory and working memory: individuals with superior memory performance also show greater preservation of task-related neural responses^156,158,159^. Here, we extend these findings to the domain of language, complementing prior evidence of the preservation of linguistic abilities and the language brain areas in aging^37,50,53,160^.

Why have some past studies found differences between younger and older adults in their brain response during language tasks? A plausible explanation for this discrepancy is that past studies have a) used paradigms that include a combination of linguistic and general cognitive demands, and b) have not isolated the *language network proper* from the nearby MD network, which gets engaged when language processing is accompanied by external task demands (e.g., ^161,162^; for reviews, see ^163,164^). For example, some have reported reduced lateralization of the language network^51^—and sometimes other networks^165^—with age, in line with the Hemispheric Asymmetry Reduction in Old Adults (HAROLD)^5^ hypothesis. In the current study, we find no difference in the degree of lateralization of the language network between younger and older adults. Because the MD network is bilateral, a task that engages both the language-selective network and the MD network—as many commonly used language paradigms do—may result in more bilateral activations in older adults because they may find the task harder and therefore rely more strongly on the MD network (see ^149^ for similar arguments in explaining past claims of more bilateral language responses in young children). Similarly, the conflation of the language network and the MD network may explain past findings of reduced functional connectivity among language-responsive areas^58,60,166^, which we do not find here (either during rest or during demanding story comprehension), when using an approach that isolates the language network from the MD network.

Another explanation for this discrepancy is that age-related neural differences during language processing only occur at higher levels of linguistic complexity (e.g., difficulty related to accessing infrequent words or processing non-local syntactic dependencies). Addressing this specific question was not the main goal of the present study. However, we did examine neural responses while participants listened to a story which included linguistically complex materials. The various linguistic complexity phenomena included in this story—such as infrequent words, unexpected words, and non-local dependencies—have been shown to engage the language network, and not the MD network, in younger adults (e.g.,^132,134^). Because investigating the effects of linguistic complexity was not the core goal, we would not want to overinterpret things absent additional data. Nevertheless, as previously mentioned, one cohort of older adults showed canonical signatures of language processing (stronger responses to infrequent and unpredictable words) in the language network. Furthermore, the language and MD networks remained clearly segregated during this story (showing ∼zero inter-network functional connectivity, similar to younger adults). Nevertheless, whether linguistic difficulty per se relies on the MD network to a greater extent with age remains to be systematically investigated by explicitly manipulating this feature. Critically, to tackle these questions without confounding general cognitive load and linguistic complexity, paradigms should be used that do not include non-linguistic task demands.

An additional reason for why some previous studies have found age-related changes in the level of response during language processing may be because they had used production tasks (e.g., ^52,167,168^). In contrast to comprehension, some aspects of language production (specifically, word retrieval) exhibit age-related decline^37,45,57^. However, comparing language comprehension and language production is not trivial. In contrast to language comprehension, which can be probed through passive listening/reading paradigms, as in the current study, eliciting brain responses during language production almost always requires specific task paradigms, such as confrontation picture naming or verbal fluency. Such paradigms engage not only the language areas, but also domain-general MD brain regions (e.g.,^109^), which deteriorate with age^5,7–13^—a finding we robustly replicate here. Some studies have reported a neural dissociation in the effect of aging on language comprehension (stable across age) versus production (a reduced response with age)^28,37^. However, the comprehension and production tasks were not matched for task demands; as a result, without independent functional localizers for the language and the MD networks, the effects are difficult to interpret: in particular, does the difference reflect a dissociation among different aspects of language processing, or does it simply reflect age-related changes in the MD network, which is engaged more strongly by the production compared to the comprehension paradigms? It is worth noting that when the language areas are identified functionally within individuals, the levels of response are strongly correlated across tasks: for example, if a participant shows a strong response during a sentence-comprehension task, they will also show a strong response during a sentence-production task (see **Supplementary Figure 2b** for evidence based on data from ^109^). Of course, these findings come from YA, and whether these correlations hold in older individuals—and therefore, whether our findings of similar response magnitudes during a comprehension task would generalize to production—remains to be established.

An important practical bottom line is that to meaningfully interpret brain responses to language—including in studies of cognitive aging—it is critical to examine the language and the MD areas separately, to understand their respective contributions. Such studies can also help resolve debates about the reasons for age-related difficulties in word retrieval, where some postulate language-specific explanations (e.g., changes in phonological encoding, ^169^), but others attribute these effects to general executive function deficits^170,171^.

Finally, to broaden the scope of our analyses beyond the core left fronto-temporal language network, in exploratory analyses (**Supplementary Figure 5**), we examined responses to language in our younger and older adults in the extended set of language-responsive areas, which include areas along the cortical midline and on the ventral temporal surface—what some refer to as the ‘extended language network’^127^. Mirroring the findings for the core language areas, most language-responsive areas in this extended network show highly similar profiles between the age groups. However, the OA group showed a lower response to language in the bilateral medial anterior SFG, and an increased response to language in the right Crus I/II/VIIb. Because these differences were not predicted a priori, we treat these results as preliminary and leave their replication and interpretation to future studies.

### The changes in the Multiple Demand network in older brains

One of the most replicated findings in the cognitive-behavioral aging literature is age-related decay in executive abilities: older adults exhibit slower processing speed, a reduced working memory capacity, and greater difficulty inhibiting distracting information^26,46^. Lower behavioral performance of older adults on the spatial working memory task in the current study aligns with these findings (**Supplementary Figure 1b**). To better understand the neural basis of these age-related changes in executive abilities, we examined the Multiple Demand (MD) network, associated with executive functions ^62^. This network is engaged by diverse working memory and cognitive control tasks^63,117,119,120^, and certain forms of reasoning, such as mathematical reasoning or understanding computer code ^172–174^. Age-related functional changes in this network have been reported in past studies^13,175^, but the effects have not been consistent^176^. For example, some have reported increased activation in older adults (sometimes referred to as a ‘compensatory increase’)^5,177^. Others have instead found an increase only at lower cognitive loads^178,179^, and suggested that at higher cognitive loads, the older adults’ neural activity may reach a plateau, leading to reduced activation compared to the younger cohort^178^. This latter proposal, where the direction of activation differences between older and younger adults depends on task difficulty is known as the Compensation-Related Utilization of Neural Circuits Hypothesis (CRUNCH; ^178^). In our study, consistent with the CRUNCH hypothesis, the response magnitude during the more difficult condition of the spatial working memory task was reduced in older adults. However, in contrast to the CRUNCH hypothesis—and in spite of a clear behavioral difference in task performance—the magnitude of response for the easier condition was similar between the age groups (if anything, the response was slightly lower, rather than higher, in older adults, partially contradicting CRUNCH).

An earlier investigation of the relationship between the MD network’s activity and behavioral performance for the spatial working memory task that was used here found that in young adults, the size of the hard > easy effect in the MD network was positively associated with task performance^126^. In particular, individuals with a larger hard > easy effect exhibited higher accuracies (and faster reaction times) for the working memory task and higher intelligence quotient (IQ) scores as measured with an independent test. Our findings of a smaller hard > easy effect in older adults (due to the lower magnitude of neural response in the hard condition; **Figure 2E**) and worse performance on the spatial WM task are consistent with these earlier findings. This concordance suggests that the same underlying factors may explain a) inter-individual differences in executive abilities within an age group and b) differences between age groups. It also aligns with studies that have shown that functional activity and connectivity patterns in the MD network are better predicted by performance on working memory tasks than by chronological age^37,180,181^. That said, we found that even when controlling for task performance, age still had an effect on response magnitudes (**Supplementary Table 3.2.5**).

### No evidence for de-differentiation: Persistent dissociation between the language and the MD networks in older adults

A prominent hypothesis that has been put forward to explain age-related cognitive decline is the de-differentiation hypothesis^182,183^, whereby areas and networks that are distinct at a younger age become less selective for a specific cognitive function and less segregated in patterns of functional connectivity in older brains (for a review, see ^184^). In contrast to compensation-based proposals, such as HAROLD and CRUNCH, which posit over-recruitment of certain brain areas (often bilaterally) under high task demands, de-differentiation concerns an intrinsic loss of selectivity and segregation, which should manifest across task/cognitive states. Much evidence for de-differentiation comes from non-human animal studies in sensorimotor cortices (e.g., ^185–187^). In humans, evidence for de-differentiation comes from fMRI studies where some have reported a) reduced selectivity of brain regions for their preferred stimulus/task and/or increased responses to some stimulus/task in regions that previously did not respond to that stimulus/task^12,183^, as well as b) reduced within-network and/or increased between-network functional connectivity^14–17,22,25^. We do not find support for the de-differentiation hypothesis in the functional response profiles of the networks investigated in this study. The language areas of older adults respond as strongly and selectively during language comprehension as those of younger adults. The selectivity holds both relative to the control condition (nonword processing), which is similar perceptually to the sentence comprehension condition, and relative to a non-linguistic working memory task. We also do not see any evidence of responses to language in the MD network of older adults, similar to young adults. These results suggest that the MD network is not recruited as an additional neural resource to process language in older individuals, at least during the kind of passive sentence comprehension paradigm used here.

With respect to the functional connectivity data, we see similarly strong within-network connectivity for the language network in the older and younger groups, and reduced within-network connectivity in the MD network, which mirrors all other neural changes in these networks. For the functional connectivity between the language and the MD areas, the results differ between rest and naturalistic story listening. During the story listening paradigm, the between-network functional connectivity is similarly low between younger and older adults. However, during resting state, we see a slight increase in connectivity between the language and the MD areas in older adults. In our understanding of the de-differentiation hypothesis, the effects of reduced within-network and increased between-network connectivity should be robust across different cognitive states, so it is unclear how to interpret these inconsistent, state-dependent effects. Minimally, the fact that the language network does not show increased connectivity with the MD network during story comprehension rules out the possibility of compensatory recruitment for language comprehension. Furthermore, another recent study has found preserved segregation of the language network from other networks at rest^57^, suggesting that these differences at rest may be small and variable across different aging cohorts. Thus, de-differentiation does not seem to ubiquitously characterize an aging brain across all networks and cognitive states. Advocates of this hypothesis should therefore offer more specific predictions about the networks for which reduced selectivity and increased integration should be observed, and under which circumstances.

Overall, different patterns of age-related change in the language versus the MD networks challenge the compensation hypotheses, whereby activation in bilateral frontal areas should increase with age across tasks^5,178^. Instead, we observed similar responses in older and younger adults in the frontal language areas during language processing, and a reduced response in the MD areas of older adults. Our results also pose a challenge to the de-differentiation hypothesis whereby networks lose their specificity with age or become more integrated with other networks^12,182,183^. We found a strong dissociation between the language and the MD networks in both the response profiles and patterns of functional connectivity in older adults, even during comprehension of a linguistically complex story. Our results are most consistent with the brain maintenance hypothesis, whereby brain networks that support functions that decline with age (e.g., executive functions) show age-related neural changes, but networks that support functions that are well-preserved in aging (e.g., language comprehension) remain young-like neurally.

### Limitations

One limitation of the current study is that we used a passive comprehension paradigm as our main language task. The topographic and functional connectivity measures are independent of the particular localizer paradigm (see **Supplementary Figure 2a**). However, the activation measures, such as the magnitude and extent of neural activity, are somewhat task-dependent (see **Supplementary Figure 2b**), so generalizing these findings to other language paradigms, including production paradigms and paradigms that systematically manipulate linguistic complexity while minimizing non-linguistic task demands, remains an important goal. The use of a passive comprehension paradigm also means that we lack a direct measure of comprehension for this task. This concern is somewhat ameliorated by the fact that during the naturalistic story listening task, which uses quite complex linguistic materials ^154,188^, our OA participants exhibited near-perfect comprehension (97% mean accuracy) and classic neural signatures of language processing (i.e., sensitivity to word frequency and surprisal) in the language network. Nevertheless, future studies should aim to relate behavioral measures of linguistic ability and neural measures more directly. In addition, measures of general cognitive abilities were only available for the OA2 cohort (where they were within the normal range)—a concern that is ameliorated by the striking degree of similarity in the neural findings between the two OA cohorts.

Another limitation is that, similar to most brain imaging work on aging (cf. ^189^), this is a cross-sectional study, and we may therefore be missing complex non-linear changes across the lifespan whose detection would require a longitudinal approach, or more dense sampling along the age continuum, along with matching the samples for factors such as educational level, occupation complexity, and specific generational experiences.

Finally, although our suite of neural measures revealed preservation of the language network with age, it remains possible that other measures will in the future uncover age-related changes in this network, including graph-theoretic measures of network organization^15^. Our exploratory analyses reveal potential age-related differences in some parts of the extended language network. We leave it to future studies to replicate these differences and understand their functional significance. Further, age differences may emerge in the time-course of information processing^34,190,191^ or in the finer-grained representations of linguistic information. Probing these differences will require temporally-sensitive and multivariate approaches.

### Conclusion

In conclusion, this study demonstrates that aging affects distinct brain networks differently, with neural changes mirroring the preservation or decline of the cognitive functions they support. Whereas the domain-general Multiple Demand network exhibits age-related decline across diverse neural measures, mirroring the decline in executive functions, the language-selective network—which supports cognitive abilities that are well-preserved in older adults—remains remarkably stable with age. These results underscore the importance of individual-level functional localization in characterizing age-related differences among adjacent networks, and highlight the need for more nuanced models of brain aging that consider the distinct trajectories of different brain networks and their associated cognitive functions.

## Materials and Methods

### Participants

509 participants were recruited from MIT and the greater Boston community. All participants were native English speakers, had normal or corrected-to-normal vision and hearing, and did not report any psychiatric or neurological disorder. These participants were recruited between 2015 and 2021 and divided into two cohorts: 483 young adults (YA, age range: 17-39; average=23.8, SD=4.5; 278 females; 431 right-handed) and 26 older adults (OA1, age range: 41-72; average=53.5, SD=8.8; 14 females; 23 right-handed). Another cohort of 38 older adults was recruited between 2021 and 2023 (OA2, age range: 44-80; average=63.9, SD=10.3; 22 females; 36 right-handed). No statistical differences were found between groups in sex ratio or handedness (*ps*>0.05). Information regarding the level of education of the participants was not available for this study. All participants gave informed consent in accordance with the requirements of MIT’s Committee on the Use of Humans as Experimental Subjects (COUHES) and were paid for their participation.

### Experimental design

Each participant completed two localizer tasks designed to identify and measure the response of the two networks of interest: a reading task for the language network and a spatial working memory task for the domain-general multiple-demand (MD) network. The language localizer task included sentences and lists of pronounceable nonwords that participants had to passively read in a blocked design, one word or nonword at a time. A simple button-press task was included at the end of each trial, to help participants remain alert. The materials are available at https://www.evlab.mit.edu/resources. The sentences > nonwords contrast targets brain regions that support high-level linguistic processing, including lexico-semantic, combinatorial syntactic, and semantic processes^79,102,121^. This task has been shown to be robust to the materials, task, and modality of presentation^73,112,192,193^. A slightly different version of the task was used for 2 of the 38 participants in OA2 where participants read sentences and nonwords (similar to the main version) but also lists of words and “Jabberwocky” sentences (morphologically and syntactically intact sentences made up of nonwords). At the end of each trial, participants had to decide whether a probe word/nonword had appeared in the immediately preceding stimulus. The sentences > nonwords contrast has been shown, across numerous studies, to not engage the MD regions, which respond more strongly during the nonwords condition^63,161,163^. Sample stimuli and trial timing details are presented in **Figure 1, Supplementary Figure 1a, and Supplementary Table 1a**. Each participant completed two runs, with condition order counterbalanced across runs.

In the Multiple Demand localizer task, participants had to keep track of four (easy condition) or eight (hard condition) sequentially presented locations in a 3 × 4 grid^63^. In both conditions, participants performed a two-alternative forced-choice task at the end of each trial to indicate the set of locations they just saw. The hard > easy contrast has been previously shown to robustly activate MD regions^63,103^, which also have been shown to respond to difficulty manipulations across many diverse tasks^63,117,118^. Participants in the YA and OA1 cohorts did one version of this task (**Figure 1**), and participants in the OA2 cohort did a slightly different version (**Supplementary Figure 1b**); minor differences in the timing/procedure do not appear to affect the activations as we had seen in cases of direct within-individual comparisons (unpublished data from the Fedorenko lab). Sample stimuli and trial timing details are presented in **Figure 1, Supplementary Figure 1a, and Supplementary Table 1b**. Each participant completed two runs, with condition order counterbalanced across runs.

A subset of the younger adults (YA) (N=82) and a subset of the older adult cohorts (OA1 N=16, OA2 N=37) also completed a resting state scan to examine functional correlations in neural activity among the language regions, among the MD regions, and between the language and the MD networks. The same YA subset and the OA2 cohort completed a second naturalistic cognition paradigm, where they passively listened to a ∼5 min-long story. The YA subset listened to a story extracted from the fairy tale *Alice in Wonderland* and OA2 listened to an edited version of the publicly available story “Elvis Died at the Florida Barber College” (by Roger Dean Kiser; unedited version: www.eastoftheweb.com/short-stories/UBooks/ElvDie.shtml). These stories were revised to incorporate various linguistic features known to increase local processing difficulty in many previous behavioral studies on sentence processing. Description of the stories can be found in^103,122,154^. The OA2 cohort responded to seven simple yes/no questions at the end of the scan across two runs to confirm their attention and comprehension of the story.

### MRI data acquisition

#### YA and OA1 cohorts

Structural and functional data were collected on the whole-body, 3 Tesla, Siemens Trio scanner with a 32-channel head coil, at the Athinoula A. Martinos Imaging Center at the McGovern Institute for Brain Research at MIT. T1-weighted structural images were collected in 176 sagittal slices with 1 mm isotropic voxels (TR = 2,530 ms, TE = 3.48 ms). Functional, blood oxygenation level dependent (BOLD), data were acquired using an EPI sequence (with a 90 degree flip angle and using GRAPPA with an acceleration factor of 2), with the following acquisition parameters: 31 4.4 mm thick near-axial slices acquired in the interleaved order (with 10% distance factor), 2.1 mm x 2.1 mm in-plane resolution, FoV in the phase encoding (A >> P) direction 200 mm and matrix size 96 mm x 96 mm, TR = 2,000 ms and TE = 30 ms. The first 10 s of each run were excluded to allow for steady state magnetization.

#### OA2 cohort

Structural and functional data were collected on a whole-body 3 Tesla Siemens Prisma scanner with a 32-channel head coil at the Athinoula A. Martinos Imaging Center at the McGovern Institute for Brain Research at MIT. T1-weighted, Magnetization Prepared Rapid Gradient Echo (MP-RAGE) structural images were collected in 176 sagittal slices with 1 mm isotropic voxels (TR = 2,530 ms, TE1 = 1.69 ms, TE2 = 3.55 ms, TE3 = 5.41 ms, TE4 = 7.27ms, TI = 1100 ms, flip = 7 degrees). Functional, blood oxygenation level-dependent (BOLD) data were acquired using an SMS EPI sequence with a 80° flip angle and using a slice acceleration factor of 3, with the following acquisition parameters: eighty-one 1.8 mm thick slices acquired in the interleaved order (with 0% distance factor), 2.4 mm × 2.4 mm in-plane resolution, FoV in the phase encoding (A >> P) direction 216 mm and matrix size 216 × 146, TR = 2,000 ms, TE = 32 ms. The first 10 s of each run were excluded to allow for steady-state magnetization. The MIT MRI team performed several systematic comparisons between the original Trio system and the system after the Prisma upgrade, and found that when preprocessing is standardized (as in the current study), the two scanners produce near-identical results.

### fMRI Preprocessing

fMRI data were analyzed using SPM12 (release 7487), CONN EvLab module (release 19b), and other custom MATLAB scripts (version 2019b). Each participant’s functional and structural data were converted from DICOM to NIFTI format. All functional scans were coregistered and resampled using B-spline interpolation to the first scan of the first session^194^. Potential outlier scans were identified from the resulting subject-motion estimates as well as from BOLD signal indicators using default thresholds in CONN preprocessing pipeline (5 standard deviations above the mean in global BOLD signal change, or framewise displacement values above 0.9 mm^195^. Functional and structural data were independently normalized into a common space (the Montreal Neurological Institute [MNI] template; IXI549Space) using SPM12 unified segmentation and normalization procedure^196^ with a reference functional image computed as the mean functional data after realignment across all timepoints omitting outlier scans. The output data were resampled to a common bounding box between MNI-space coordinates (-90, -126, -72) and (90, 90, 108), using 2 mm isotropic voxels and 4th order spline interpolation for the functional data, and 1mm isotropic voxels and trilinear interpolation for the structural data. Last, the functional data were smoothed spatially with a 4 mm FWHM Gaussian kernel. Data from the two naturalistic paradigms (resting-state and story listening) were further preprocessed by regressing out of each voxel’s time-course principal components of the six subject-specific motion parameters and BOLD signal time-courses extracted from the white matter and cerebrospinal fluid. Residuals were then bandpass filtered (0.0100-0.2500 Hz).

### First-level analysis

Responses in individual voxels were estimated using a General Linear Model (GLM) in which each experimental condition was modeled with a boxcar function convolved with the canonical hemodynamic response function (HRF) (fixation was modeled implicitly, such that all timepoints that did not correspond to one of the conditions were assumed to correspond to a fixation period). Temporal autocorrelations in the BOLD signal timeseries were accounted for by a combination of high-pass filtering with a 128 seconds cutoff, and whitening using an AR(0.2) model (first-order autoregressive model linearized around the coefficient α=0.2) to approximate the observed covariance of the functional data in the context of Restricted Maximum Likelihood estimation (ReML). In addition to experimental condition effects, the GLM design included first-order temporal derivatives for each condition (included to model variability in the HRF delays), as well as nuisance regressors to control for the effect of slow linear drifts, subject-specific motion parameters (6 parameters), and potential outlier scans (identified during preprocessing as described above) on the BOLD signal.

### Functional localization of the language and MD networks and response estimation

For each participant, functional regions of interest (fROIs) were defined using the Group-constrained Subject-Specific (GSS) approach^73^, whereby a set of parcels or “search spaces” (i.e., brain areas within which most individuals in prior studies showed activity for the localizer contrast) is combined with each individual participant’s activation map for the same or similar contrast. To define the language fROIs, we used five parcels derived from a group-level representation of data for the sentences > nonwords contrast in 220 independent participants. For each participant, within each parcel, the top 10% most active voxels were selected based on the t values for the sentences > nonwords contrast of the language task. These parcels have been used in much prior work (e.g., ^122,197–199^) and include three regions in the left frontal cortex: two located in the inferior frontal gyrus (LH IFG and LH IFGorb), and one located in the middle frontal gyrus (LH MFG); and two regions in the left temporal cortex spanning the entire extent of the lateral temporal lobe (LH AntTemp and LH PostTemp). Additionally, we examined activations in the right hemisphere (RH) homotopes of the language regions. To define the fROIs in the RH, the left hemisphere parcels were mirror-projected onto the RH to create five homotopic parcels. By design, the parcels cover relatively large swaths of cortex in order to be able to accommodate inter-individual variability. Hence the mirrored versions are likely to encompass RH language regions despite possible hemispheric asymmetries in the precise locations of activations (for validation, see ^76,103,192,200^). In addition, in exploratory analyses, we used an extended set of language parcels created based on 706 participants who all completed a reading-based language localizer task^105^. In addition to the core LH and RH frontal and temporal areas, this set of parcels includes areas along the cortical midline, in the temporal pole, on the ventral temporal surface, and in the cerebellum (the parcels are available at https://osf.io/7594t/overview;^127^).

To define the MD fROIs, we used a set of 20 parcels (10 in each hemisphere) derived from a group-level representation of data for the hard > easy spatial working memory contrast in 197 participants. For each participant, within each parcel, the top 10% most active voxels were selected based on the t values for the hard > easy spatial WM contrast. These parcels have been used in prior work^122,173^ and include regions in the posterior, middle, and anterior parietal cortex (LH and RH postParietal, midParietal, and antParietal), in the superior frontal gyrus (LH and RH supFrontal), in the precentral gyrus (LH and RH PrecG), in the IFG pars opercularis (LH and RH IFGop), in the middle frontal gyrus and its orbital part (LH and RH midFront and MidFrontOrb), in the insula (LH and RH insula), and in the medial frontal cortex (LH and RH medialFront).

For both the language and the MD networks, we estimated the responses of these individually defined fROIs to the sentences and nonwords conditions of the language task, and the easy and hard conditions of the MD task. For extracting the responses of the language fROIs to the language task conditions, and for extracting the responses of the MD fROIs to the spatial WM task conditions, an across-runs cross-validation procedure was used^81^, ensuring independence of data^201^.

To ensure that the spatial topography of the language network did not depend on the particular localizer used, we used an approach from Shain & Fedorenko^125^. In particular, for a subset of 431 young adults and 53 older adults (OA1 = 31, OA2 = 23), we identified the language network from each individual’s functional connectome (based on functional correlations computed across diverse fMRI tasks; for more details on the methodology of individualized functional connectomics, see ^125^). We then computed a correlation between each individual’s unthresholded, grey-matter masked whole-brain language localizer activation map and their probabilistic estimate of the language network derived from the connectivity data. We then tested whether these correlations change with age using a linear model with continuous age or age group as fixed effects. The results are shown in **Supplementary Figure 2a**.

### Topography measures

To assess individual variability in the spatial organization of the language and MD networks, we computed two complementary metrics for each participant. First, to measure global activation pattern similarity, we computed the spatial correlation (SpCorr) between each participant’s unthresholded whole-brain activation map (Sentences>Nonwords contrast map for the language network, Hard>Easy contrast map for the MD network) and previously published standard probabilistic atlas for the respective network created based on statistical maps thresholded at the top 10% most active voxels for the relevant contrast (language: n=806, MD: n=691, both available for download from https://www.evlab.mit.edu/resources). To create a relative index of typicality, each participant’s SpCorr value was then mean-centered by subtracting the average SpCorr of the entire study sample for that network (note that results were highly similar when mean-centering at the group level). Second, to measure regional displacement of peak activation, we calculated the Euclidean distance between individual-specific activation peaks and group-level probabilistic atlas peak. For each functional language parcel (or “search space”) a group-level peak was defined as a maximum value from a probabilistic language or MD atlas in each parcel (using the same atlases as for SpCorr). Individual-specific activation peaks were defined within the same functional parcels as a voxel with the maximum t-value for the relevant contrast map. The Euclidean distance (in mm) between these two points in MNI space was then calculated, with a larger distance indicating greater spatial deviation of an individual’s functional activation hotspot from the canonical group location.

### Measures of activation extent, lateralization, and spatial overlap

Each participant’s activation map for each localizer contrast (sentences > nonwords for language and hard > easy spatial working memory for MD) were thresholded at an alpha level 0.05 after FDR correction and binarized. To determine activation extent, the number of significant voxels was computed at the whole-brain level and within each language and MD parcel. To determine the lateralization index (which was only done for the language network, as the MD network does not show a strong hemispheric bias), the number of contrast-activated voxels in the right hemisphere (RH) at the FDR-corrected *p*<0.05 significance threshold was subtracted from the number of contrast-activated voxels in the left hemisphere (LH) at the same threshold, and the resulting value was divided by the sum of contrast-activated voxels across hemispheres: (LH-RH)/(LH+RH). Finally, Dice overlap coefficients ^141^ were first computed between the FDR-corrected activation maps of the two language runs and between the FDR-corrected activation maps of the two MD runs to measure within-network overlap. To measure the overlap between the two networks, a Dice coefficient was computed for each language-MD pair of runs and then averaged across the four pairs.

### Functional connectivity measures

For each participant, we first averaged BOLD timeseries across voxels within each individual-specific language and MD fROI (i.e., top 10% voxels most responsive to the language and MD localizer tasks). Fisher-transformed Pearson correlations were then estimated between averaged timeseries of pairs of language and MD fROIs for each naturalistic paradigm (i.e., resting-state and story listening) for each participant to measure within and between-network connectivity.

### Sensitivity to frequency and surprisal in the older adults’ language network’s response during naturalistic story comprehension

We modeled responses to each word during the naturalistic story comprehension task as a function of frequency and surprisal, with controls for word rate and end-of-sentence markers. Corpus frequency was computed using a KenLM unigram model trained on Gigaword 3. Surprisal was estimated using GPT2-xl^133^. A deconvolutional intercept was used for word rate, corresponding to a vector of ones time-aligned with the word onsets of the audio stimuli. Rate captures influences of stimulus timing independently of stimulus properties (see e.g., ^202,203^). All predictors were convolved with the canonical hemodynamic response function prior to regression, and used to fit an ordinary least squares (OLS) model that jointly used rate, end-of-sentence, frequency, and surprisal to predict average BOLD within each language fROI and within the language network as a whole, with independent OLS models for each participant. Group-level effects were estimated and tested using a summary statistics approach (a t-test of the average effect estimate across the group within each fROI and in the network as a whole).

### Statistical analyses

All statistical analyses were performed in R (version 4.2.1). To provide a comprehensive assessment of age-related differences in the function of the language and MD networks, we conducted a series of statistical analyses using a combination of linear mixed-effects models (LMMs) using the *lmer* function as part of the *lme4* package^204^, beta regressions, and ANOVAs. Across all analyses, we tested the effects of age using three parallel approaches: 1) a two-level categorical factor comparing Younger Adults (YA) vs. Older Adults (OA), 2) a three-level categorical factor comparing YA vs. each OA cohort separately (OA1, OA2) to assess any cohort-specific effects, and 3) a mean-centered continuous age factor to assess any linear trends not captured by categorical group comparisons. Model assumptions for the final models were confirmed through visual inspection of residual plots. P-values from all analyses were corrected for multiple comparisons, as detailed below, with a significance threshold of α=0.05.

#### 1. Examining age-related differences in behavior, topography, and activation-related measures for the networks of interest

##### Behavioral performance

MD task accuracy was first compared across cohorts using a linear mixed-effects model to investigate the effects of cohort and task condition on accuracy, as well as their interaction. The model included cohort, condition, and their interaction as fixed effects, with a random intercept for participant (Accuracy ∼ Cohort*Condition + (1|Participant)). To further investigate the differences in accuracy between groups within each condition, we conducted post-hoc pairwise comparisons using Tukey-adjusted estimated marginal means to control for multiple comparisons.

##### Topography

To examine age-related differences in functional network topography, we tested the effects of age on two complementary metrics: global similarity and regional peak deviation. First, to assess global activation pattern similarity, we fit LMMs predicting the relative typicality index (mean-centered spatial correlation). Separate models tested the interaction between network atlas (Language vs. MD) and either a categorical age group factor or a continuous age factor. These models included a random intercept for participant. Post-hoc tests using estimated marginal means (*emmeans*) were then conducted to assess group differences within each network. Second, to assess regional peak deviation, we analyzed the Euclidean distance from probabilistic atlas peaks. These LMMs tested the interaction between network and age (both categorical and continuous). To account for multiple sources of non-independence, these models included crossed random intercepts for participant and parcel. For the language network, this analysis was restricted to left-hemisphere parcels. To probe significant interactions, follow-up models were run to examine the simple linear effect of age within each network separately.

##### Activation extent

To examine age-related differences in the spatial extent of activation, the voxel count (thresholded at FDR-corrected *p*<0.05) was first square-root transformed to stabilize variance. At the aggregated network and hemisphere levels, we used a Likelihood Ratio Test (LRT)-informed LMM approach to select between homoscedastic (*lmer*) and heteroscedastic (*lme*) models for categorical predictors and further account for potential differences in group variance. Models included a fixed effect for cohort or mean-centered age and random intercepts for participant. A random intercept for parcel was also included when the voxel count was measured within broad anatomical parcels (Eq. 1a-b). At the exploratory individual ROI level, simpler linear models were used. P-values for main effects from aggregated-level models were corrected using the Holm-Bonferroni method. At the ROI-level, p-values were FDR-corrected at α=0.05 across all ROIs for each combination of task and age predictor.

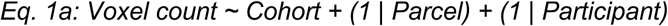

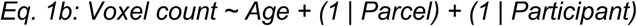

##### Lateralization

One-way ANOVAs (using *aov_ez* function of the *afex* package) were used to test for differences in lateralization of the language network activity among the categorical age groups (2-level and 3-level). The effect of continuous age was assessed using a simple linear regression.

##### Response magnitude

To evaluate 1) the response of the language and MD networks to both the language task (i.e., the size of the sentences > nonwords effect) and the MD task (i.e., the size of the hard > easy effect) within cohort and 2) age-related differences across cohorts, the response magnitude for each task condition was first transformed using a Yeo-Johnson power transformation to better satisfy model assumptions. The optimal transformation parameter (λ) was estimated separately for each analysis using the *powerTransform* function from the *car* package. All analyses were performed at the network, hemisphere, and fROI levels.

###### Identification of task effects within each group

At the network and hemisphere levels, within-group task effects were evaluated with LMMs including a fixed effect for task condition and a random intercept for participant and fROI (Eq. 2). P-value approximation was performed with the *lmerTest* package^205^. Multiple comparison corrections (Holm-Bonferrroni for network and hemisphere levels, FDR for fROI level) were applied separately within each age group.

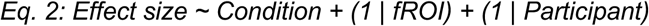

###### Age-related effects on BOLD response magnitude

To evaluate age-related differences in BOLD response magnitude in each network, analyses were tailored to the predictor type and analysis level. Models evaluating differences in response magnitude included a fixed effect for the interaction between cohort or mean-centered age and task condition. For the categorical age predictors at the aggregated (network/hemisphere) levels, we used a data-driven approach. An LRT determined if modeling heteroscedasticity provided a superior fit. If significant (*p*<0.05), a heteroscedastic LMM was fit using *nlme::lme* with variance structure allowing group-specific residual variances (varIdent weights).

Otherwise, a homoscedastic LMM was fit using *lme4::lmer*. All models used a crossed random-effects structure for participants and fROIs (Eq. 3a). For the continuous age predictor at the aggregated levels, a homoscedastic LMM (*lme4::lmer*) was used with the same crossed random-effects structure (Eq. 3b). For all three predictor types at the exploratory fROI level, a homoscedastic LMM (*lme4::lmer*) with a random intercept for participant was fit for each ROI. P-values for the primary interaction terms at the aggregated levels were corrected across all within- and between-network tests using the Holm-Bonferroni method.

For the ROI-level analyses, we controlled the FDR at *p*<0.05 across all ROIs within each specific analysis (e.g., all ROIs testing the S>N effect for the YA vs. OA comparison).

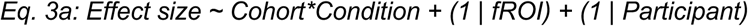

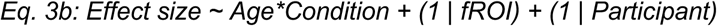

###### Post-hoc brain-behavior analysis

To explore the relationship between task performance and brain activity, we conducted a post-hoc analysis for the spatial working memory (MD) task. We fit a set of LMMs predicting response magnitude from the three-way interaction between task condition (Hard vs. Easy), age (categorical or continuous), and task accuracy (mean-centered). This allowed us to determine whether age-related differences in brain activity were modulated by individual differences in performance. An LRT was again used to select between homoscedastic and heteroscedastic models for the categorical age predictors. P-values for the three-way interaction terms were corrected using the Holm-Bonferroni method.

##### Spatial overlap

First, to evaluate whether the spatial overlap within (language-language, MD-MD) and between (language-MD) domains changed with age, we analyzed Dice coefficients between activation maps from pairs of BOLD runs thresholded at FDR-corrected *p*<0.05 for the relevant task contrast. Because Dice coefficients are bounded between 0 and 1, we used beta regression (*betareg* package), which is specifically designed for such proportional data. To handle potential values of 0 or 1, the Dice coefficients were first transformed using the standard (*y* * (*n*-1) + 0.5) / *n* formula where *n* is the sample size^206^. We tested for age-related effects across the three types of age predictors (2-level group, 3-level group, continuous) on spatial overlap within and between language and MD task activation maps. Multiple comparisons across tasks were corrected using the Holm-Bonferroni method.

Next, to directly compare differences in overlap between domain types within each age group, we implemented an LMM (*lmer*). This model predicted the transformed Dice coefficient from the full interaction between domain type (within-Language, within-MD, between language and MD) and age group (tested with both 2- and 3-level group factors), including a random intercept for participant. Following this model, we conducted planned post-hoc contrasts using estimated marginal means (*emmeans*) to test, within each age group, whether within-network overlap differed significantly from between-network overlap. P-values from contrasts were also adjusted using Holm-Bonferroni correction.

### 2. Examining age-related differences in inter-regional correlations within and between the language and the MD networks

To assess age-related differences in functional connectivity, we analyzed Fisher-transformed inter-regional correlations within and between networks during resting-state and story listening. Similar to the response magnitude analyses, we used an LRT-informed LMM approach to test the effects of our three age predictors (2-level group, 3-level group, continuous) at the hemisphere and network levels. Models included a fixed effect for cohort or age and random intercepts for fROI pair and participant (e.g., Eq. 4 for categorical model). Multiple comparisons across paradigms (i.e., resting-state and story listening) and analysis levels (network, hemisphere) were corrected using the Holm-Bonferroni method.

In addition, to directly compare connectivity strength across network types, we implemented an LMM (*lmer*) for each task paradigm. These models predicted Fisher-transformed correlation coefficients from the interaction between connectivity type (within-Language, within-MD, or between-network) and age group (2-level and 3-level group factors). All models included crossed random intercepts for subject and for each fROI-pair (Eq. 5). Following each model, we performed planned post-hoc contrasts using estimated marginal means (*emmeans*) to specifically test whether within-network connectivity (both within-Language and within-MD) differed from between-network connectivity, conducted separately for each age group. P-values from all resulting contrasts were corrected for multiple comparisons using a single Holm-Bonferroni procedure.

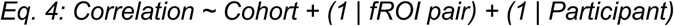

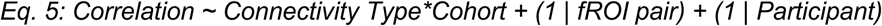

## Data availability

The data that support the findings of this study will be available on the paper website (https://osf.io/2q65t/). Source Data are provided with this paper.

## Code availability

The code that supports the findings of this study will be available via the paper website (https://osf.io/2q65t/).

## Acknowledgments

We acknowledge the Athinoula A. Martinos Imaging Center at the McGovern Institute for Brain Research, MIT, including the technical support team (Steve Shannon and Atsushi Takahashi). We thank all individuals who participated in this study. We additionally acknowledge present and past members of the Boston University Center for Brain Recovery and the Fedorenko Lab at MIT. This work was supported by the NIH - National Institute on Deafness and Other Communication Disorders (grant R01DC016950 to SK and EF). EF was additionally supported by research funds from the McGovern Institute for Brain Research, MIT’s Simons Center for the Social Brain, MIT’s Poitras Center for Psychiatric Disorders Research, and MIT’s Quest for Intelligence.

## Notes

https://osf.io/2q65t/?view_only=ab1833db12c64eb0a7cc61c5795d35cd

### Competing Interest Statement

Dr. Kiran is a scientific advisor for Constant Therapy Health, but there is no overlap between this role and the submitted investigation. The other authors report no conflicts.

### Summary of Updates

The main changes from the first submission are as follows: i) Strengthened statistical framework for analyzing age-related differences: We have combined the older adults' data across the two cohorts and now present a simpler two-group comparison (Younger Adults vs. all Older Adults), supplemented with a continuous model of age to capture linear age-related trajectories across the adult lifespan. Our original comparisons between the YA group and each of the older adult cohorts (YA vs. OA1, and YA vs. OA2) are included in the Supplementary Information. For all categorical analyses, we now formally test for and model heteroscedasticity (unequal variance between groups), ensuring greater statistical validity. All the critical results hold in this new analytic framework. ii) We performed an analysis that shows that the network defined with the language localizer corresponds closely to a network that can be recovered with no task contrast, solely from patterns of functional correlations (Supp. Figures 2a-b). iii) We have included two new analyses to evaluate the stability of the activation topography of the language network (and our control, MD network) in aging. The first analysis measures the global similarity between an individual's whole-brain activation pattern and a normative probabilistic atlas for the same network. The second analysis focuses on finer-grained, more focal age-related shifts by measuring the spatial deviation in the locations of individual activation peaks from the normative-atlas centroids within the regions of interest for each network. iv) We analyzed comprehension accuracy for the questions asked after the story comprehension task; this analysis revealed that accuracies were near ceiling, in spite of the fact that the story used complex linguistic materials. We also tested whether OA's responses during the naturalistic story condition showed signatures of linguistic complexity effects that have been previously observed in younger adults. Similar to past studies in younger adults, we found that OA showed strong sensitivity to frequency and surprisal. Together these new analyses suggest that comprehension was intact in OA and qualitatively similar to YA (at least for basic measures of linguistic comprehension). v) In exploratory analyses, we have tested whether age-related activation changes occur in a broader set of language-sensitive brain regions, which together comprise the "extended language network" (Wolna et al., 2025). This analysis (included in Supp. Fig. 5) offers a more comprehensive overview of aging effects beyond the core language regions that we focus on in our main analyses.

