## Supplementary for "The language network ages well: Preserved topography, lateralization, selectivity, and within-network functional connectivity in older brains"

### TABLE OF CONTENTS

**1. Additional details of the language and Multiple Demand (MD) localizers.**
- Supp. Table 1a. Details of the language localizer tasks.
- Supp. Table 1b. Details of the MD localizer tasks.
- Supp. Figure 1a. Alternative versions of the language and MD localizers, used for small subsets of participants.
- Supp. Figure 1b. Behavioral performance on the MD spatial working memory task across cohorts.

**2. The language network's topography does not depend on a particular localizer and its response magnitude in individual participants varies in similar ways across paradigms.**
- Supp. Figure 2a. The language network as defined by the current language localizer corresponds well with the language network derived from voxel-wise patterns of functional connectivity in both young and older adults.
- Supp. Figure 2b. Correlations in the magnitude of the language network's response across four language comprehension and production tasks.

**3. Details and outputs of the statistical models for the main results**
- **Supp. Tables 3.1. The language network is well preserved in aging.**
  - 3.1.1.Spatial correlations with the language network atlas
  - 3.1.2.Deviations from the peak locations in the language network atlas
  - 3.1.3.Lateralization
  - 3.1.4.Activation Extent
  - 3.1.5.Response Magnitude
  - 3.1.6.Functional Connectivity
- **Supp. Tables 3.2. The Multiple Demand network shows pronounced age-related changes.**
  - 3.2.1.Spatial correlations with the MD network atlas
  - 3.2.2.Deviations from the peak locations in the MD network atlas
  - 3.2.3.Activation Extent
  - 3.2.4.Response Magnitude
  - 3.2.5.Response magnitude, controlling for task performance
  - 3.2.6.Functional Connectivity
- **Supp. Tables 3.3. The language and the MD networks remain robustly segregated in older adults.**
  - 3.3.1.Response Magnitude to the non-preferred domain
  - 3.3.2.Activation Overlap
  - 3.3.3.Functional Connectivity

**4. Key results broken down by fROI and OA cohort.**
(complementary to main **Figure 2**)
- Supp. Figure 4a. Activation extent of the language and the MD contrasts in the younger adults (YA) and older adults(OA1 and OA2).
- Supp. Figure 4b. Response profiles of the language fROIs and MD fROIs in younger adults (YA) and older adults (OA1 and OA2).

**5. Exploratory analyses of the Extended Language Network**
- Supp. Figure 5. Response profiles of the non-canonical language fROIs in younger and older adults.

### 6. Details of the statistical models for additional results

- **Supp. Tables 6.1. Response magnitude results.**
  - 6.1.1. Network level: Language response in Language fROIs (LH + RH)
  - 6.1.2. Network level: Language response in MD fROIs (LH + RH)
  - 6.1.3. Network level: MD response in Language fROIs (LH+RH)
  - 1.2.5. Hemisphere level: MD response in the MD LH fROIs
  - 1.2.6. Hemisphere level: MD response in the MD RH fROIs
- **Supp. Tables 6.2. Activation extent results.**
  - **6.2.1. Extent of Whole-Brain Activation Maps (FDR $p$<0.05)**
    - 6.2.1.1. Language Contrast S>N
    - 6.2.1.2. MD Contrast: H>E
  - 6.2.2. Extent of Activation Maps within Parcels (FDR $p$<0.05)
    - 6.2.2.1. Language Contrast: S>N (LH+RH)
    - 6.2.2.2. MD Contrast: H>E (LH)
    - 6.2.2.3. MD Contrast: H>E (RH)
- **Supp. Tables 6.3. Functional connectivity results.**
  - 6.3.1. Resting-State: Within Language Network
    - 6.3.1.1. Network Level (All Connections)
    - 6.3.1.2. Hemisphere Level (Inter-hemispheric)
  - **6.3.2. Story Listening: Within Language Network**
    - 6.3.2.1. Network Level (All Connections)
    - 6.3.2.2. Hemisphere Level (Inter-hemispheric)
  - **6.3.3. Resting-State: Within MD Network**
    - 6.3.3.1. Network Level (All Connections)
    - 6.3.3.2. Hemisphere Level (Inter-hemispheric)
  - **6.3.4. Story Listening: Within MD Network**
    - 6.3.4.1. Network Level (All Connections)
    - 6.3.4.2. Hemisphere Level (Inter-hemispheric)
  - **6.3.5. Resting-State: Between Language and MD Networks**
    - 6.3.5.1. Network Level (All Connections)
    - 6.3.5.2. Hemisphere Level (Inter-hemispheric)
  - **6.3.6. Story Listening: Between Language and MD Networks**
    - 6.3.6.1. Network Level (All Connections)
    - 6.3.6.2. Hemisphere Level (Inter-hemispheric)
- **Supp. Tables 6.4. Spatial activation overlap results within domain between runs**
  - 6.4.1. Within Language Network
  - 6.4.2. Within MD Network

### 1. Additional details of the language and Multiple Demand (MD) localizers.

| Version | Main | Alternate |
|---|---|---|
| Number of participants | 483 (YA), 26 (OA1), 36 (OA2) | 2 (OA2) |
| Task type | Button press | Memory probe |
| Words/ Nonwords per trial | 12 | 8 |
| Trial duration (ms) | 6000 | 6000 |
| Trial-initial interval (ms) | 100 | 0 |
| Stimulus (ms) | 5400 (450/word) | 3600 (450/word) |
| Pre-probe interval (ms) | 0 | 400 |
| Button icon/ Memory probe (ms) | 400 | 1500 |
| Trial-final interval (ms) | 100 | 500 |
| Trials per block | 3 | 3 |
| Block duration (s) | 18 | 18 |
| Blocks per condition per run | 8 | 6 |
| Conditions | Sentences, Nonwords | Sentences, Wordlists, Jabberwocky, Nonwords |
| Fixation block duration (s) | 14 | 12 |
| Fixation blocks per run | 5 | 7 |
| Total run time (s) | 358 | 516 |

**Supp. Table 1a. Details of language localizer tasks.**

| Version | Main | Alternate |
|---|---|---|
| Number of participants | 483 (YA), 26 (OA1) | 38 (OA2) |
| Trial-initial fixation (ms) | 500 | 500 |
| Stimulus (ms) | 4000 (1000/square flash) | 4000 (1000/square flash) |
| Time for choice condition | 3000 (max, response triggers feedback) | 3500 |
| Feedback duration (ms) | 250 | 0 (not provided) |
| Post-feedback fixation (ms) | 3250 – Reaction Time | 0 |
| Trial length (s) | 8 | 8 |
| Trials per block | 4 | 3 |
| Block duration (s) | 32 | 24 |
| Blocks per condition per run | 6 | 6 |
| Conditions | Hard, Easy | Hard, Easy |
| Fixation block duration (s) | 16 | 12 |
| Fixation blocks per run | 4 | 4 |
| Total run time (s) | 448 | 336 |

**Supp. Table 1b. Details of MD localizer tasks.**

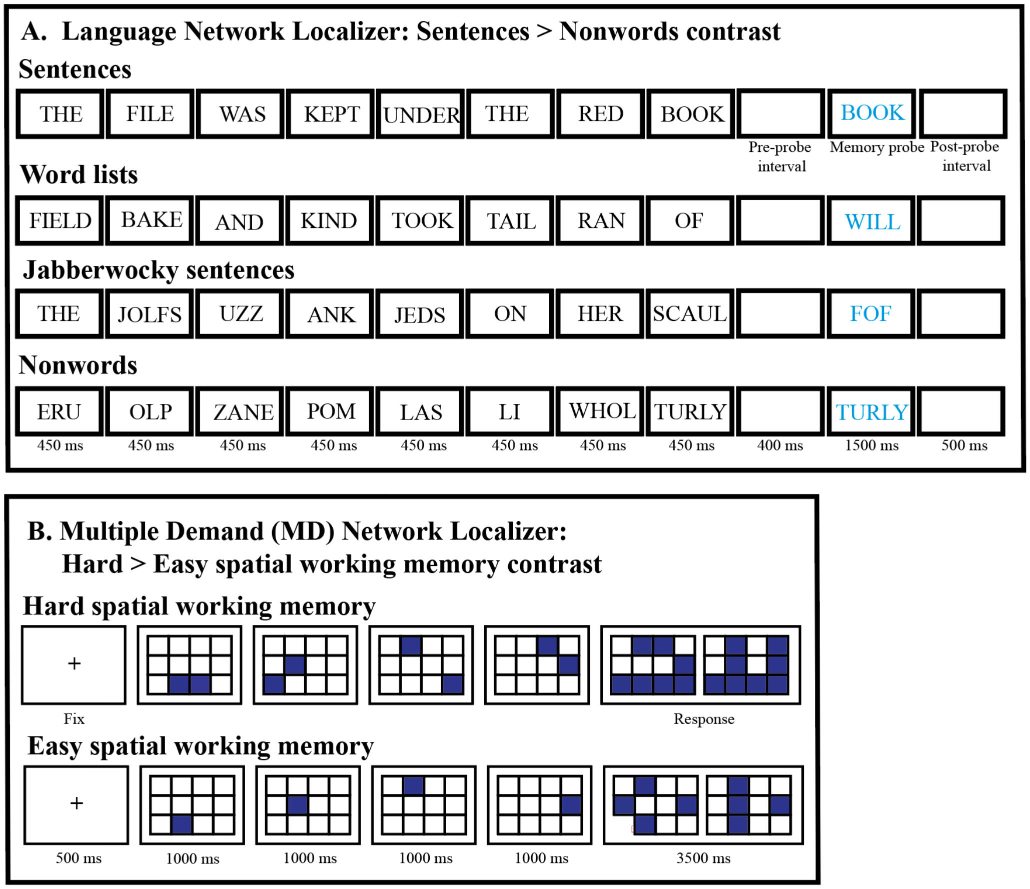


**Supp. Figure 1a. Alternative versions of the language and MD localizers, used for subsets of participants (see Materials and Methods).** The main versions are displayed in Figure 1 in the main paper.

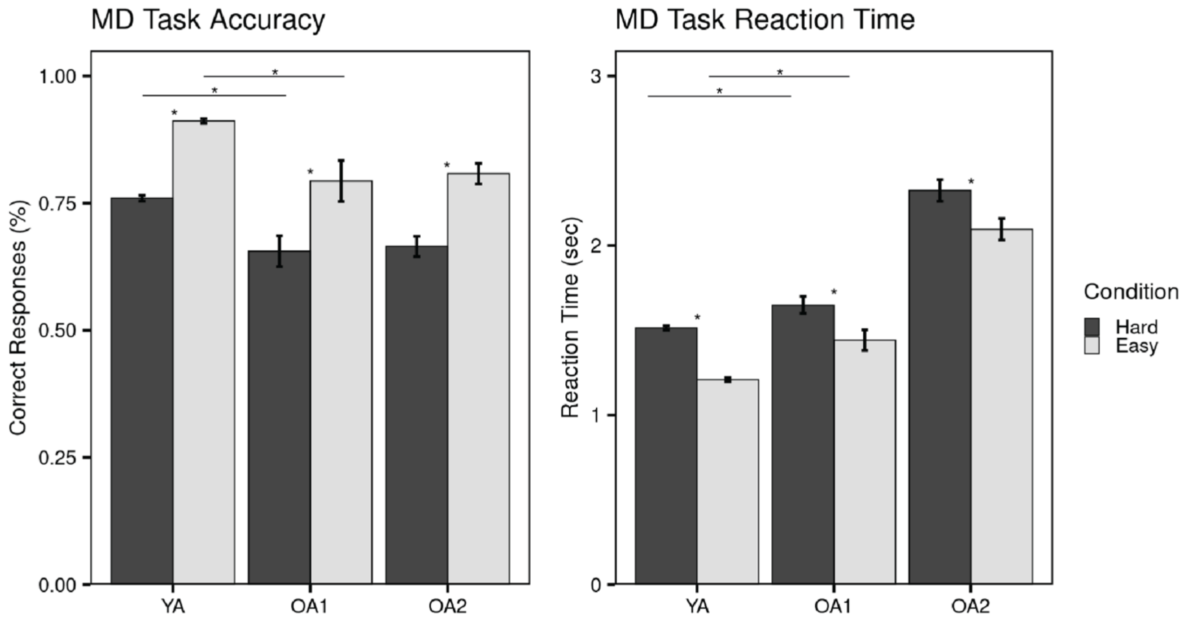


**Supp. Figure 1b. Behavioral performance on the Multiple Demand spatial working memory task across cohorts.** Note that accuracy and reaction time are not directly comparable between YA and OA2, given that the trials were structured differently. As a result, when discussing the behavioral data, we focus on the accuracy comparison between YA and OA1. Asterisks represent significant differences between conditions within groups ($p<0.05$), asterisks above a line represent significant differences between groups ($p<0.05$).

### 2. The language network's topography does not depend on a particular localizer and its response magnitude in individual participants varies in similar ways across paradigms.

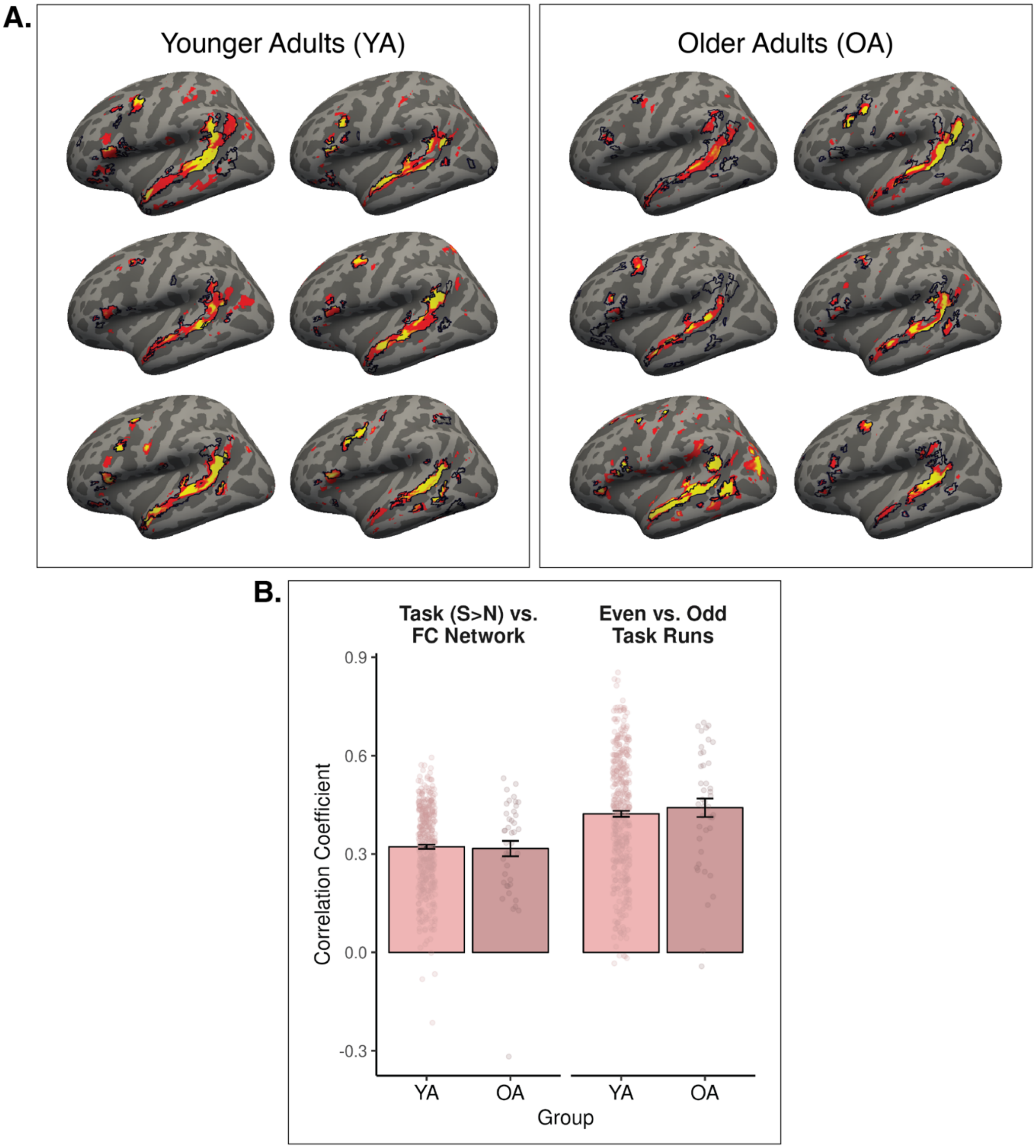


**Supp. Figure 2a. The language network as defined by the current language localizer corresponds well with the language network derived from voxel-wise patterns of functional connectivity in both young and older adults.** A/ Correspondence between localizer contrast maps (colored, yellow = stronger contrast magnitude / red = weaker contrast magnitude) and maps derived from patterns of functional connectivity (black outlines) for sample participants. B/ Average spatial correlations between the language localizer contrast maps (Sentences > Nonwords) and maps derived from patterns of functional connectivity (left), and, for comparison, between two language localizer task runs.

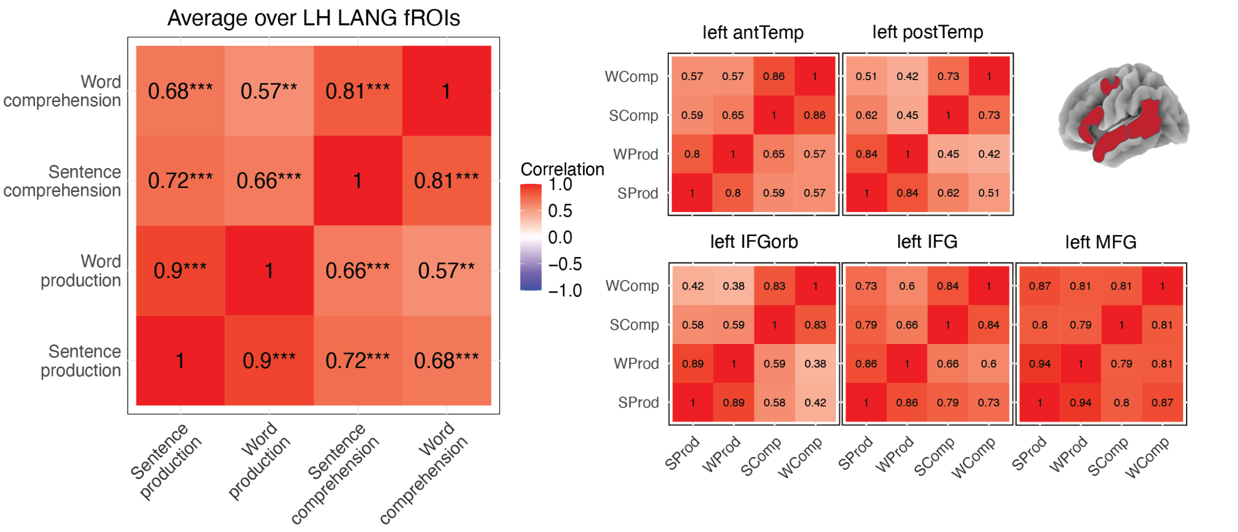


**Supp. Figure 2b. Correlations in the magnitude of the language network's response across four language comprehension and production tasks.** To illustrate that the language network response magnitudes are correlated across different language task paradigms, we leveraged a previously published fMRI dataset (Hu, Small et al., 2023) which included 29 young adults performing four language tasks: sentence comprehension, word comprehension, sentence production, and word production. The language network's response to these four tasks showed strong correlations across participants.

### 3. Details and outputs of the statistical models for the main results.

#### Supp. Tables 3.1. The language network is well preserved in aging.

##### 3.1.1.Spatial correlations with the language network atlas

| Atlas | Predictor | Estimate | Std. Error | Statistic | 95% CI | p-value | p.adj |
|---|---|---|---|---|---|---|---|
| Language | (Intercept) | 0.000 | 0.005 | 0.00 | (-0.009, 0.009) | 0.999 | |
| | Age | -0.000 | 0.000 | -0.32 | (-0.001, 0.001) | 0.746 | 0.746 |

| Atlas | Predictor | Estimate | Std. Error | Statistic | 95% CI | p-value | p.adj |
|---|---|---|---|---|---|---|---|
| Language | (Intercept) | 0.002 | 0.005 | 0.44 | (-0.007, 0.012) | 0.659 | |
| | Group = OA | -0.019 | 0.014 | -1.29 | (-0.047, 0.010) | 0.198 | 0.198 |

| Atlas | Predictor | Estimate | Std. Error | Statistic | 95% CI | p-value | p.adj |
|---|---|---|---|---|---|---|---|
| Language | (Intercept) | 0.002 | 0.005 | 0.44 | (-0.007, 0.012) | 0.659 | |
| | Group = OA1 | -0.041 | 0.022 | -1.87 | (-0.083, 0.002) | 0.062 | 0.124 |
| | Group = OA2 | -0.003 | 0.018 | -0.19 | (-0.039, 0.032) | 0.852 | 0.852 |

##### 3.1.2.Deviations from the peak locations in the language network atlas

| Atlas | Predictor | Estimate | Std. Error | Statistic | 95% CI | p-value | p.adj |
|---|---|---|---|---|---|---|---|
| Language | (Intercept) | 10.338 | 1.148 | 9.00 | (7.152, 13.524) | <.001 | |
| | Age | -0.014 | 0.007 | -1.97 | (-0.028, -0.000) | 0.049 | **0.049** |

| Atlas | Predictor | Estimate | Std. Error | Statistic | 95% CI | p-value | p.adj |
|---|---|---|---|---|---|---|---|
| Language | (Intercept) | 10.406 | 1.149 | 9.06 | (7.220, 13.591) | <.001 | |
| | Group = OA | -0.585 | 0.288 | -2.03 | (-1.150, -0.019) | 0.043 | **0.043** |

| Atlas | Predictor | Estimate | Std. Error | Statistic | 95% CI | p-value | p.adj |
|---|---|---|---|---|---|---|---|
| Language | (Intercept) | 10.406 | 1.149 | 9.06 | (7.220, 13.591) | <.001 | |
| | Group = OA1 | -0.801 | 0.441 | -1.82 | (-1.667, 0.065) | 0.070 | 0.139 |
| | Group = OA2 | -0.442 | 0.362 | -1.22 | (-1.154, 0.269) | 0.223 | 0.223 |

##### 3.1.3.Lateralization

| Predictor | Estimate | Std. Error | Statistic | 95% CI (Lower) | 95% CI (Upper) | p-value |
|---|---|---|---|---|---|---|
| (Intercept) | 0.530 | 0.014 | 39.173 | 0.504 | 0.557 | <.001 |
| Age | -0.001 | 0.001 | -0.538 | -0.003 | 0.002 | 0.591 |

| Predictor | Estimate | Std. Error | Statistic | 95% CI (Lower) | 95% CI (Upper) | p-value |
|---|---|---|---|---|---|---|
| (Intercept) | 0.535 | 0.014 | 37.175 | 0.507 | 0.563 | <.001 |
| Group = OA | -0.041 | 0.042 | -0.973 | -0.124 | 0.042 | 0.331 |

| Predictor | Estimate | Std. Error | Statistic | 95% CI (Lower) | 95% CI (Upper) | p-value |
|---|---|---|---|---|---|---|
| (Intercept) | 0.535 | 0.014 | 37.152 | 0.507 | 0.563 | <.001 |

| Predictor | Estimate | Std. Error | Statistic | 95% CI (Lower) | 95% CI (Upper) | p-value |
|---|---|---|---|---|---|---|
| Group = OA1 | -0.067 | 0.063 | -1.059 | -0.192 | 0.057 | 0.290 |
| Group = OA2 | -0.023 | 0.054 | -0.422 | -0.128 | 0.083 | 0.673 |

#### 3.1.4.Activation Extent

**Left Hemisphere Language Parcels**

| Predictor | Estimate | Std. Error | Statistic | 95% CI (Lower) | 95% CI (Upper) | p-value | p.adj |
|---|---|---|---|---|---|---|---|
| (Intercept) | 16.256 | 3.743 | 4.343 | 5.900 | 26.613 | 0.012 | |
| Group = OA1 | -0.407 | 1.167 | -0.349 | -2.700 | 1.886 | 0.727 | 1.000 |
| Group = OA2 | 1.298 | 0.977 | 1.329 | -0.621 | 3.217 | 0.185 | 0.738 |

| Predictor | Estimate | Std. Error | Statistic | 95% CI (Lower) | 95% CI (Upper) | p-value | p.adj |
|---|---|---|---|---|---|---|---|
| (Intercept) | 16.326 | 3.742 | 4.363 | 5.968 | 26.684 | 0.012 | |
| Age | 0.022 | 0.019 | 1.152 | -0.016 | 0.060 | 0.250 | 0.315 |

**Right Hemisphere Language Parcels**

| Predictor | Estimate | Std. Error | Statistic | 95% CI (Lower) | 95% CI (Upper) | p-value | p.adj |
|---|---|---|---|---|---|---|---|
| (Intercept) | 8.274 | 2.485 | 3.329 | 1.427 | 15.120 | 0.028 | |
| Group = OA1 | 0.858 | 1.150 | 0.746 | -1.402 | 3.118 | 0.456 | 1.000 |
| Group = OA2 | 1.581 | 0.963 | 1.642 | -0.311 | 3.472 | 0.101 | 0.607 |

| Predictor | Estimate | Std. Error | Statistic | 95% CI (Lower) | 95% CI (Upper) | p-value | p.adj |
|---|---|---|---|---|---|---|---|
| (Intercept) | 8.423 | 2.483 | 3.392 | 1.574 | 15.272 | 0.027 | |
| Age | 0.031 | 0.019 | 1.623 | -0.006 | 0.068 | 0.105 | 0.315 |

#### 3.1.5. Response Magnitude

**Language response in the Language LH fROIs**

| Predictor | Estimate | Std. Error | Statistic | 95% CI (Lower) | 95% CI (Upper) | p-value | p.adj |
|---|---|---|---|---|---|---|---|
| (Intercept) | 0.435 | 0.015 | 28.079 | 0.404 | 0.465 | <.001 | |
| Condition = S | 0.754 | 0.008 | 91.523 | 0.738 | 0.770 | <.001 | |
| Group = OA1 | -0.076 | 0.069 | -1.104 | -0.211 | 0.059 | 0.270 | |
| Group = OA2 | -0.070 | 0.060 | -1.154 | -0.188 | 0.049 | 0.249 | |
| (Condition = S) x (Group = OA1) | 0.004 | 0.037 | 0.115 | -0.069 | 0.077 | 0.909 | 1 |
| (Condition = S) x (Group = OA2) | 0.135 | 0.041 | 3.294 | 0.054 | 0.215 | 0.001 | **0.005** |

| Predictor | Estimate | Std. Error | Statistic | 95% CI (Lower) | 95% CI (Upper) | p-value | p.adj |
|---|---|---|---|---|---|---|---|
| (Intercept) | 0.426 | 0.113 | 3.780 | 0.117 | 0.736 | 0.018 | |
| Condition = S | 0.763 | 0.010 | 75.713 | 0.744 | 0.783 | 0.000 | |
| Age | -0.002 | 0.001 | -2.096 | -0.005 | 0.000 | 0.036 | |
| (Condition = S) x Age | 0.003 | 0.001 | 3.401 | 0.001 | 0.004 | 0.001 | **0.002** |

**Language response in the Language RH fROIs**

| Predictor | Estimate | Std. Error | Statistic | 95% CI (Lower) | 95% CI (Upper) | p-value | p.adj |
|---|---|---|---|---|---|---|---|
| (Intercept) | 0.254 | 0.017 | 14.877 | 0.221 | 0.288 | <.001 | |

| | | | | | | | |
|---|---|---|---|---|---|---|---|
| Condition = S | 0.373 | 0.008 | 44.781 | 0.357 | 0.390 | <.001 | |
| Group = OA1 | -0.024 | 0.074 | -0.317 | -0.169 | 0.122 | 0.751 | |
| Group = OA2 | 0.038 | 0.064 | 0.593 | -0.088 | 0.163 | 0.554 | |
| (Condition = S) x (Group = OA1) | 0.006 | 0.030 | 0.195 | -0.053 | 0.065 | 0.846 | 1 |
| (Condition = S) x (Group = OA2) | 0.060 | 0.033 | 1.828 | -0.004 | 0.124 | 0.068 | 0.271 |

| Predictor | Estimate | Std. Error | Statistic | 95% CI (Lower) | 95% CI (Upper) | p-value | p.adj |
|---|---|---|---|---|---|---|---|
| (Intercept) | 0.256 | 0.044 | 5.862 | 0.145 | 0.367 | 0.002 | |
| Condition = S | 0.378 | 0.010 | 37.649 | 0.358 | 0.397 | <.001 | |
| Age | 0.000 | 0.001 | -0.007 | -0.003 | 0.002 | 0.994 | |
| (Condition = S) x Age | 0.001 | 0.001 | 1.239 | -0.001 | 0.003 | 0.215 | 0.215 |

#### 3.1.6.Functional Connectivity

**Left Hemisphere language fROIs – Resting State**

| Predictor | Estimate | Std. Error | Statistic | 95% CI (Lower) | 95% CI (Upper) | p-value | p.adj |
|---|---|---|---|---|---|---|---|
| (Intercept) | 0.566 | 0.044 | 12.738 | 0.470 | 0.663 | <.001 | |
| Group = OA1 | -0.012 | 0.044 | -0.279 | -0.099 | 0.075 | 0.781 | 1.000 |
| Group = OA2 | 0.024 | 0.032 | 0.750 | -0.039 | 0.087 | 0.455 | 1.000 |

| Predictor | Estimate | Std. Error | Statistic | 95% CI (Lower) | 95% CI (Upper) | p-value | p.adj |
|---|---|---|---|---|---|---|---|
| (Intercept) | 0.571 | 0.043 | 13.282 | 0.477 | 0.666 | <.001 | |
| Age | 0.000 | 0.001 | 0.392 | -0.001 | 0.002 | 0.696 | 0.696 |

**Left Hemisphere language fROIs – Story Listening**

| Predictor | Estimate | Std. Error | Statistic | 95% CI (Lower) | 95% CI (Upper) | p-value | p.adj |
|---|---|---|---|---|---|---|---|
| (Intercept) | 0.768 | 0.053 | 14.513 | 0.654 | 0.882 | <.001 | |
| Group = OA2 | -0.008 | 0.040 | -0.210 | -0.088 | 0.071 | 0.834 | 1.000 |

| Predictor | Estimate | Std. Error | Statistic | 95% CI (Lower) | 95% CI (Upper) | p-value | p.adj |
|---|---|---|---|---|---|---|---|
| (Intercept) | 0.765 | 0.051 | 14.899 | 0.653 | 0.877 | <.001 | |
| Age | 0.000 | 0.001 | -0.254 | -0.002 | 0.002 | 0.800 | 1.000 |

**Right Hemisphere language fROIs – Resting State**

| Predictor | Estimate | Std. Error | Statistic | 95% CI (Lower) | 95% CI (Upper) | p-value | p.adj |
|---|---|---|---|---|---|---|---|
| (Intercept) | 0.357 | 0.044 | 8.145 | 0.262 | 0.453 | <.001 | |
| Group = OA1 | 0.006 | 0.041 | 0.151 | -0.075 | 0.087 | 0.880 | 1.000 |
| Group = OA2 | 0.056 | 0.030 | 1.861 | -0.003 | 0.115 | 0.065 | 0.390 |

| Predictor | Estimate | Std. Error | Statistic | 95% CI (Lower) | 95% CI (Upper) | p-value | p.adj |
|---|---|---|---|---|---|---|---|
| (Intercept) | 0.374 | 0.043 | 8.760 | 0.279 | 0.468 | <.001 | |
| Age | 0.001 | 0.001 | 1.057 | -0.001 | 0.002 | 0.292 | 0.585 |

**Right Hemisphere language fROIs – Story Listening**

| Predictor | Estimate | Std. Error | Statistic | 95% CI (Lower) | 95% CI (Upper) | p-value | p.adj |
|---|---|---|---|---|---|---|---|
| (Intercept) | 0.526 | 0.060 | 8.794 | 0.396 | 0.656 | <.001 | |
| Group = OA2 | 0.020 | 0.042 | 0.469 | -0.063 | 0.103 | 0.640 | 1.000 |

| Predictor | Estimate | Std. Error | Statistic | 95% CI (Lower) | 95% CI (Upper) | p-value | p.adj |
|---|---|---|---|---|---|---|---|
| (Intercept) | 0.532 | 0.058 | 9.125 | 0.404 | 0.660 | <.001 | |
| Age | 0.000 | 0.001 | -0.170 | -0.002 | 0.002 | 0.865 | 1.000 |

**Supp. Tables 3.2. The Multiple Demand network shows pronounced age-related changes.**

**3.2.1. Spatial correlations with the MD network atlas**

| Atlas | Predictor | Estimate | Std. Error | Statistic | 95% CI | p-value | p.adj |
|---|---|---|---|---|---|---|---|
| MD | (Intercept) | -0.000 | 0.006 | -0.02 | (-0.012, 0.012) | 0.984 | |
| | Age | -0.005 | 0.000 | -9.97 | (-0.006, -0.004) | < 0.001 | **<.001** |

| Atlas | Predictor | Estimate | Std. Error | Statistic | 95% CI | p-value | p.adj |
|---|---|---|---|---|---|---|---|
| MD | (Intercept) | 0.021 | 0.007 | 3.12 | (0.008, 0.034) | 0.002 | |
| | Group = OA | -0.182 | 0.020 | -9.16 | (-0.221, -0.143) | < 0.001 | **<.001** |

| Atlas | Predictor | Estimate | Std. Error | Statistic | 95% CI | p-value | p.adj |
|---|---|---|---|---|---|---|---|
| MD | (Intercept) | 0.021 | 0.007 | 3.11 | (0.008, 0.034) | 0.002 | |
| | Group = OA1 | -0.190 | 0.030 | -6.22 | (-0.249, -0.130) | < 0.001 | **<.001** |
| | Group = OA2 | -0.177 | 0.025 | -7.08 | (-0.226, -0.128) | < 0.001 | **<.001** |

**3.2.2. Deviations from the peak locations in the MD network atlas**

| Atlas | Predictor | Estimate | Std. Error | Statistic | 95% CI | p-value | p.adj |
|---|---|---|---|---|---|---|---|
| MD | (Intercept) | 11.405 | 0.371 | 30.78 | (10.632, 12.179) | < 0.001 | |
| | Age | 0.035 | 0.006 | 6.23 | (0.024, 0.046) | < 0.001 | **<.001** |

| Atlas | Predictor | Estimate | Std. Error | Statistic | 95% CI | p-value | p.adj |
|---|---|---|---|---|---|---|---|
| MD | (Intercept) | 11.241 | 0.371 | 30.26 | (10.466, 12.016) | < 0.001 | |
| | Group = OA | 1.416 | 0.223 | 6.35 | (0.978, 1.854) | < 0.001 | **<.001** |

| Atlas | Predictor | Estimate | Std. Error | Statistic | 95% CI | p-value | p.adj |
|---|---|---|---|---|---|---|---|
| MD | (Intercept) | 11.241 | 0.371 | 30.26 | (10.466, 12.016) | < 0.001 | |
| | Group = OA1 | 1.312 | 0.342 | 3.84 | (0.641, 1.983) | < 0.001 | **<.001** |
| | Group = OA2 | 1.485 | 0.281 | 5.29 | (0.934, 2.036) | < 0.001 | **<.001** |

**3.2.3. Activation Extent**

**Bilateral**

| Predictor | Estimate | Std. Error | Statistic | 95% CI (Lower) | 95% CI (Upper) | p-value | p.adj |
|---|---|---|---|---|---|---|---|
| (Intercept) | 19.218 | 0.431 | 44.598 | 18.373 | 20.063 | <.001 | |
| Group = OA1 | -8.566 | 1.924 | -4.451 | -12.346 | -4.786 | <.001 | **<.001** |
| Group = OA2 | -9.444 | 1.586 | -5.953 | -12.560 | -6.328 | <.001 | **<.001** |

| Predictor | Estimate | Std. Error | Statistic | 95% CI (Lower) | 95% CI (Upper) | p-value | p.adj |
|---|---|---|---|---|---|---|---|
| (Intercept) | 18.163 | 1.417 | 12.821 | 15.228 | 21.097 | <.001 | |
| Age | -0.231 | 0.032 | -7.335 | -0.293 | -0.169 | <.001 | **<.001** |

#### 3.2.4. Response Magnitude

**MD response in bilateral MD fROIs**

| | Estimate | Std. Error | Statistic | 95% CI (Lower) | 95% CI (Upper) | p-value | p.adj |
|---|---|---|---|---|---|---|---|
| (Intercept) | 1.277 | 0.025 | 50.332 | 1.227 | 1.327 | <.001 | |
| Condition = H | 0.586 | 0.004 | 132.768 | 0.577 | 0.594 | <.001 | |
| Group = OA1 | -0.037 | 0.121 | -0.308 | -0.275 | 0.201 | 0.758 | |
| Group = OA2 | -0.149 | 0.094 | -1.596 | -0.333 | 0.035 | 0.111 | |
| (Condition = H) x (Group = OA1) | -0.287 | 0.019 | -15.243 | -0.324 | -0.250 | <.001 | **<.001** |
| (Condition = H) x (Group = OA2) | -0.347 | 0.015 | -22.777 | -0.377 | -0.317 | <.001 | **<.001** |

| | Estimate | Std. Error | Statistic | 95% CI (Lower) | 95% CI (Upper) | p-value | p.adj |
|---|---|---|---|---|---|---|---|
| (Intercept) | 1.265 | 0.093 | 13.569 | 1.071 | 1.458 | <.001 | |
| Condition = H | 0.549 | 0.008 | 71.629 | 0.534 | 0.564 | <.001 | |
| Age | -0.002 | 0.002 | -1.147 | -0.006 | 0.002 | 0.252 | |
| (Condition = H) x Age | -0.009 | 0.001 | -15.160 | -0.010 | -0.008 | <.001 | **<.001** |

#### 3.2.5. Response magnitude, controlling for task performance

**Bilateral MD fROIs**

| Predictor | Estimate | Std. Error | Statistic | p-value |
|---|---|---|---|---|
| (Intercept) | 1.328 | 0.028 | 46.925 | <.001 |
| Condition = Hard | 0.618 | 0.026 | 23.493 | **<.001** |
| Group = OA1 | -0.040 | 0.115 | -0.352 | 0.725 |
| Accuracy | -0.534 | 0.162 | -3.305 | **0.001** |
| (Condition = Hard) x (Group = OA1) | -0.100 | 0.116 | -0.863 | 0.388 |
| (Condition = Hard) x (Accuracy) | 1.453 | 0.153 | 9.488 | **<.001** |
| (Group=OA1) x (Accuracy) | 0.415 | 0.384 | 1.082 | 0.279 |
| (Condition = Hard) x (Group=OA1) x (Accuracy) | -0.190 | 0.489 | -0.388 | 0.698 |

| Predictor | Estimate | Std. Error | Statistic | p-value |
|---|---|---|---|---|
| (Intercept) | 1.326 | 0.095 | 13.893 | <.001 |
| Condition = Hard | 0.607 | 0.026 | 23.264 | **<.001** |
| Age | -0.002 | 0.003 | -0.618 | 0.537 |
| Accuracy | -0.507 | 0.156 | -3.247 | **0.001** |
| (Condition = Hard) x (Age) | -0.006 | 0.003 | -2.033 | **0.043** |
| (Condition = Hard) x (Accuracy) | 1.386 | 0.148 | 9.355 | **<.001** |
| (Age) x (Accuracy) | 0.011 | 0.011 | 0.946 | 0.344 |

| | | | | |
|---|---|---|---|---|
| (Condition = Hard) x (Age) x (Accuracy) | -0.013 | 0.015 | -0.869 | 0.386 |

#### 3.2.6. Functional Connectivity

**Left Hemisphere MD fROIs – Resting State**

| Predictor | Estimate | Std. Error | Statistic | 95% CI (Lower) | 95% CI (Upper) | p-value | p.adj |
|---|---|---|---|---|---|---|---|
| (Intercept) | 0.433 | 0.025 | 17.279 | 0.383 | 0.482 | <.001 | |
| Group = OA1 | -0.114 | 0.038 | -2.983 | -0.190 | -0.038 | 0.003 | 0.017 |
| Group = OA2 | -0.091 | 0.028 | -3.215 | -0.146 | -0.035 | 0.002 | **0.010** |

| Predictor | Estimate | Std. Error | Statistic | 95% CI (Lower) | 95% CI (Upper) | p-value | p.adj |
|---|---|---|---|---|---|---|---|
| (Intercept) | 0.394 | 0.023 | 17.257 | 0.349 | 0.439 | <.001 | |
| Age | -0.003 | 0.001 | -4.535 | -0.004 | -0.002 | <.001 | **<.001** |

**Right Hemisphere MD fROIs – Resting State**

| Predictor | Estimate | Std. Error | Statistic | 95% CI (Lower) | 95% CI (Upper) | p-value | p.adj |
|---|---|---|---|---|---|---|---|
| (Intercept) | 0.473 | 0.024 | 19.729 | 0.426 | 0.521 | <.001 | |
| Group = OA1 | -0.082 | 0.037 | -2.204 | -0.156 | -0.008 | 0.029 | 0.059 |
| Group = OA2 | -0.142 | 0.027 | -5.168 | -0.196 | -0.087 | <.001 | **<.001** |

| Predictor | Estimate | Std. Error | Statistic | 95% CI (Lower) | 95% CI (Upper) | p-value | p.adj |
|---|---|---|---|---|---|---|---|
| (Intercept) | 0.424 | 0.022 | 19.478 | 0.381 | 0.467 | <.001 | |
| Age | -0.004 | 0.001 | -5.937 | -0.005 | -0.002 | <.001 | **<.001** |

**Left Hemisphere MD fROIs – Story Listening**

| Predictor | Estimate | Std. Error | Statistic | 95% CI (Lower) | 95% CI (Upper) | p-value | p.adj |
|---|---|---|---|---|---|---|---|
| (Intercept) | 0.408 | 0.024 | 16.788 | 0.360 | 0.456 | <.001 | |
| Group = OA2 | -0.089 | 0.024 | -3.660 | -0.137 | -0.041 | <.001 | **0.001** |

| Predictor | Estimate | Std. Error | Statistic | 95% CI (Lower) | 95% CI (Upper) | p-value | p.adj |
|---|---|---|---|---|---|---|---|
| (Intercept) | 0.380 | 0.023 | 16.587 | 0.334 | 0.425 | <.001 | |
| Age | -0.002 | 0.001 | -4.083 | -0.004 | -0.001 | <.001 | **<.001** |

**Right Hemisphere MD fROIs – Story Listening**

| Predictor | Estimate | Std. Error | Statistic | 95% CI (Lower) | 95% CI (Upper) | p-value | p.adj |
|---|---|---|---|---|---|---|---|
| (Intercept) | 0.446 | 0.023 | 18.971 | 0.399 | 0.492 | <.001 | |
| Group = OA2 | -0.109 | 0.022 | -4.838 | -0.153 | -0.064 | <.001 | **<.001** |

| Predictor | Estimate | Std. Error | Statistic | 95% CI (Lower) | 95% CI (Upper) | p-value | p.adj |
|---|---|---|---|---|---|---|---|
| (Intercept) | 0.411 | 0.022 | 18.439 | 0.367 | 0.456 | <.001 | |
| Age | -0.003 | 0.001 | -4.909 | -0.004 | -0.002 | <.001 | **<.001** |

### Supp. Tables 3.3. The language and the MD networks remain robustly segregated in older adults.

#### 3.3.1. Response Magnitude to the non-preferred domain

**Language response in the MD LH fROIs**

| Predictor | Estimate | Std. Error | Statistic | 95% CI (Lower) | 95% CI (Upper) | p-value | p.adj |
|---|---|---|---|---|---|---|---|
| (Intercept) | 0.589 | 0.094 | 6.301 | 0.381 | 0.797 | <.001 | |
| Condition = S | -0.274 | 0.011 | -24.715 | -0.296 | -0.253 | <.001 | |
| Group = OA1 | -0.193 | 0.104 | -1.861 | -0.397 | 0.011 | 0.063 | |
| Group = OA2 | -0.135 | 0.085 | -1.580 | -0.302 | 0.033 | 0.115 | |
| (Condition = S) x (Group = OA1) | 0.073 | 0.050 | 1.451 | -0.025 | 0.171 | 0.147 | 1 |
| (Condition = S) x (Group = OA2) | 0.035 | 0.041 | 0.850 | -0.046 | 0.115 | 0.395 | 1 |

| Predictor | Estimate | Std. Error | Statistic | 95% CI (Lower) | 95% CI (Upper) | p-value | p.adj |
|---|---|---|---|---|---|---|---|
| (Intercept) | 0.571 | 0.093 | 6.130 | 0.364 | 0.779 | <.001 | |
| Condition = S | -0.269 | 0.010 | -25.742 | -0.289 | -0.248 | <.001 | |
| Age | -0.005 | 0.002 | -2.912 | -0.008 | -0.002 | 0.004 | |
| (Condition = S) x Age | 0.002 | 0.001 | 2.500 | 0.000 | 0.004 | 0.012 | 0.075 |

**Language response in the MD RH fROIs**

| Predictor | Estimate | Std. Error | Statistic | 95% CI (Lower) | 95% CI (Upper) | p-value | p.adj |
|---|---|---|---|---|---|---|---|
| (Intercept) | 0.550 | 0.082 | 6.746 | 0.370 | 0.730 | <.001 | |
| Condition = S | -0.219 | 0.010 | -21.053 | -0.240 | -0.199 | <.001 | |
| Group = OA1 | -0.130 | 0.110 | -1.185 | -0.346 | 0.085 | 0.236 | |
| Group = OA2 | 0.044 | 0.090 | 0.491 | -0.133 | 0.221 | 0.623 | |
| (Condition = S) x (Group = OA1) | 0.032 | 0.047 | 0.687 | -0.060 | 0.124 | 0.492 | 1 |
| (Condition = S) x (Group = OA2) | 0.026 | 0.039 | 0.677 | -0.049 | 0.102 | 0.498 | 1 |

| Predictor | Estimate | Std. Error | Statistic | 95% CI (Lower) | 95% CI (Upper) | p-value | p.adj |
|---|---|---|---|---|---|---|---|
| (Intercept) | 0.547 | 0.081 | 6.746 | 0.368 | 0.727 | <.001 | |
| Condition = S | -0.216 | 0.010 | -22.055 | -0.235 | -0.197 | <.001 | |
| Age | -0.001 | 0.002 | -0.524 | -0.004 | 0.003 | 0.601 | |
| (Condition = S) x Age | 0.001 | 0.001 | 1.026 | -0.001 | 0.002 | 0.305 | 1 |

**MD response in the Language LH fROIs**

| Predictor | Estimate | Std. Error | Statistic | 95% CI (Lower) | 95% CI (Upper) | p-value | p.adj |
|---|---|---|---|---|---|---|---|
| (Intercept) | -0.060 | 0.026 | -2.303 | -0.111 | -0.009 | 0.021 | |
| Condition = H | -0.163 | 0.009 | -18.098 | -0.181 | -0.146 | <.001 | |
| Group = OA1 | 0.097 | 0.116 | 0.838 | -0.130 | 0.325 | 0.402 | |
| Group = OA2 | -0.162 | 0.096 | -1.682 | -0.350 | 0.027 | 0.093 | |
| (Condition = H) x (Group = OA1) | -0.007 | 0.033 | -0.202 | -0.070 | 0.057 | 0.840 | 1 |
| (Condition = H) x (Group = OA2) | 0.036 | 0.032 | 1.118 | -0.027 | 0.100 | 0.264 | 1 |

| Predictor | Estimate | Std. Error | Statistic | 95% CI (Lower) | 95% CI (Upper) | p-value | p.adj |
|---|---|---|---|---|---|---|---|
| (Intercept) | -0.067 | 0.269 | -0.248 | -0.808 | 0.675 | 0.816 | |
| Condition = H | -0.161 | 0.018 | -9.012 | -0.196 | -0.126 | <.001 | |

| | | | | | | | |
|---|---|---|---|---|---|---|---|
| Age | -0.001 | 0.002 | -0.438 | -0.005 | 0.003 | 0.661 | |
| (Condition = H) x Age | 0.001 | 0.001 | 0.936 | -0.001 | 0.004 | 0.349 | 1 |

**MD response in the Language RH fROIs**

| Predictor | Estimate | Std. Error | Statistic | 95% CI (Lower) | 95% CI (Upper) | p-value | p.adj |
|---|---|---|---|---|---|---|---|
| (Intercept) | 0.019 | 0.025 | 0.762 | -0.030 | 0.069 | 0.446 | |
| Condition = H | -0.084 | 0.009 | -9.253 | -0.101 | -0.066 | <.001 | |
| Group = OA1 | 0.244 | 0.111 | 2.185 | 0.025 | 0.462 | 0.029 | |
| Group = OA2 | -0.028 | 0.092 | -0.307 | -0.210 | 0.153 | 0.759 | |
| (Condition = H) x (Group = OA1) | 0.042 | 0.025 | 1.665 | -0.008 | 0.092 | 0.096 | 0.959 |
| (Condition = H) x (Group = OA2) | 0.001 | 0.027 | 0.049 | -0.051 | 0.054 | 0.961 | 1 |

| Predictor | Estimate | Std. Error | Statistic | 95% CI (Lower) | 95% CI (Upper) | p-value | p.adj |
|---|---|---|---|---|---|---|---|
| (Intercept) | 0.028 | 0.212 | 0.134 | -0.554 | 0.611 | 0.900 | |
| Condition = H | -0.082 | 0.018 | -4.632 | -0.116 | -0.047 | <.001 | |
| Age | 0.003 | 0.002 | 1.350 | -0.001 | 0.006 | 0.177 | |
| (Condition = H) x Age | 0.001 | 0.001 | 0.744 | -0.002 | 0.004 | 0.457 | 1 |

#### 3.3.2.Activation Overlap between Language and MD task maps

| Predictor | Estimate | Std. Error | Statistic | 95% CI (Lower) | 95% CI (Upper) | p-value | p.adj |
|---|---|---|---|---|---|---|---|
| (Intercept) | -2.834 | 0.034 | -84.390 | -2.900 | -2.768 | <.001 | |
| Group = OA1 | 0.154 | 0.123 | 1.250 | -0.088 | 0.396 | 0.211 | 0.819 |
| Group = OA2 | -0.148 | 0.117 | -1.268 | -0.377 | 0.081 | 0.205 | 0.819 |

| Predictor | Estimate | Std. Error | Statistic | 95% CI (Lower) | 95% CI (Upper) | p-value | p.adj |
|---|---|---|---|---|---|---|---|
| (Intercept) | -2.836 | 0.032 | -88.606 | -2.898 | -2.773 | <.001 | |
| Age | -0.002 | 0.002 | -0.681 | -0.006 | 0.003 | 0.496 | 0.991 |

#### 3.3.3.Functional Connectivity between Language and MD fROIs

**Left Hemisphere – Resting State**

| Predictor | Estimate | Std. Error | Statistic | 95% CI (Lower) | 95% CI (Upper) | p-value | p.adj |
|---|---|---|---|---|---|---|---|
| (Intercept) | 0.083 | 0.019 | 4.302 | 0.045 | 0.122 | <.001 | |
| Group = OA1 | -0.016 | 0.031 | -0.525 | -0.076 | 0.044 | 0.600 | 1.000 |
| Group = OA2 | 0.089 | 0.022 | 4.005 | 0.045 | 0.133 | <.001 | **<.001** |

| Predictor | Estimate | Std. Error | Statistic | 95% CI (Lower) | 95% CI (Upper) | p-value | p.adj |
|---|---|---|---|---|---|---|---|
| (Intercept) | 0.106 | 0.018 | 5.921 | 0.071 | 0.142 | <.001 | |
| Age | 0.002 | 0.001 | 2.788 | 0.000 | 0.003 | 0.006 | **0.007** |

**Right Hemisphere – Resting State**

| Predictor | Estimate | Std. Error | Statistic | 95% CI (Lower) | 95% CI (Upper) | p-value | p.adj |
|---|---|---|---|---|---|---|---|
| (Intercept) | 0.051 | 0.019 | 2.670 | 0.013 | 0.088 | 0.009 | |
| Group = OA1 | 0.061 | 0.029 | 2.133 | 0.004 | 0.118 | 0.035 | 0.139 |

| Predictor | Estimate | Std. Error | Statistic | 95% CI (Lower) | 95% CI (Upper) | p-value | p.adj |
|---|---|---|---|---|---|---|---|
| Group = OA2 | 0.071 | 0.021 | 3.365 | 0.029 | 0.112 | <.001 | **0.005** |

| Predictor | Estimate | Std. Error | Statistic | 95% CI (Lower) | 95% CI (Upper) | p-value | p.adj |
|---|---|---|---|---|---|---|---|
| (Intercept) | 0.077 | 0.018 | 4.409 | 0.043 | 0.112 | <.001 | |
| Age | 0.001 | 0.001 | 2.985 | 0.001 | 0.002 | 0.003 | **0.007** |

**Left Hemisphere Level – Story Listening**

| Predictor | Estimate | Std. Error | Statistic | 95% CI (Lower) | 95% CI (Upper) | p-value | p.adj |
|---|---|---|---|---|---|---|---|
| (Intercept) | 0.059 | 0.021 | 2.781 | 0.017 | 0.102 | 0.006 | |
| Group = OA2 | 0.013 | 0.024 | 0.552 | -0.034 | 0.060 | 0.582 | 0.582 |

| Predictor | Estimate | Std. Error | Statistic | 95% CI (Lower) | 95% CI (Upper) | p-value | p.adj |
|---|---|---|---|---|---|---|---|
| (Intercept) | 0.064 | 0.020 | 3.179 | 0.024 | 0.103 | 0.002 | |
| Age | 0.000 | 0.001 | 0.722 | -0.001 | 0.002 | 0.472 | 0.472 |

**Right Hemisphere – Story Listening**

| Predictor | Estimate | Std. Error | Statistic | 95% CI (Lower) | 95% CI (Upper) | p-value | p.adj |
|---|---|---|---|---|---|---|---|
| (Intercept) | 0.044 | 0.020 | 2.196 | 0.004 | 0.083 | 0.030 | |
| Group = OA2 | 0.039 | 0.022 | 1.795 | -0.004 | 0.082 | 0.075 | 0.166 |

| Predictor | Estimate | Std. Error | Statistic | 95% CI (Lower) | 95% CI (Upper) | p-value | p.adj |
|---|---|---|---|---|---|---|---|
| (Intercept) | 0.056 | 0.019 | 3.003 | 0.019 | 0.093 | 0.003 | |
| Age | 0.001 | 0.001 | 1.604 | 0.000 | 0.002 | 0.111 | 0.229 |

**<u>4. Key results broken down by fROI and OA cohort</u>** <u>(complementary to main Figure 2).</u>

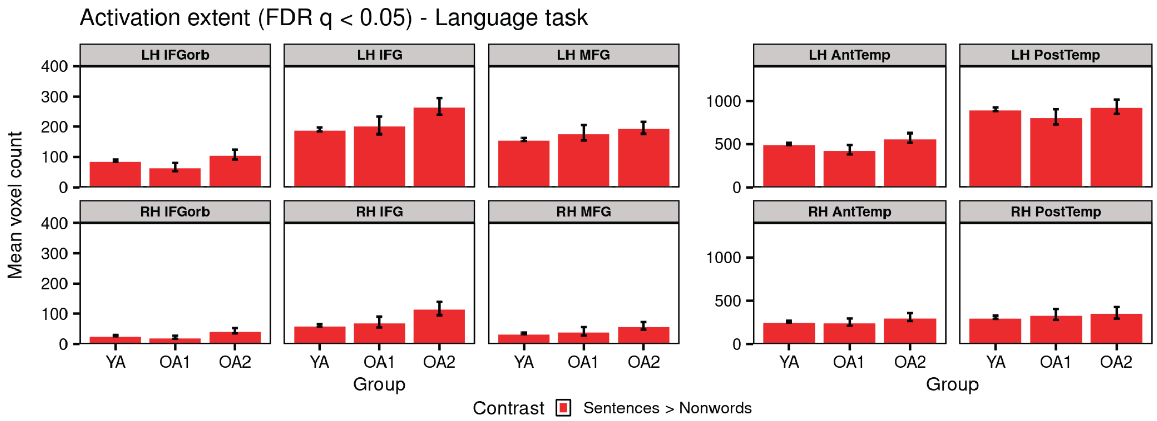


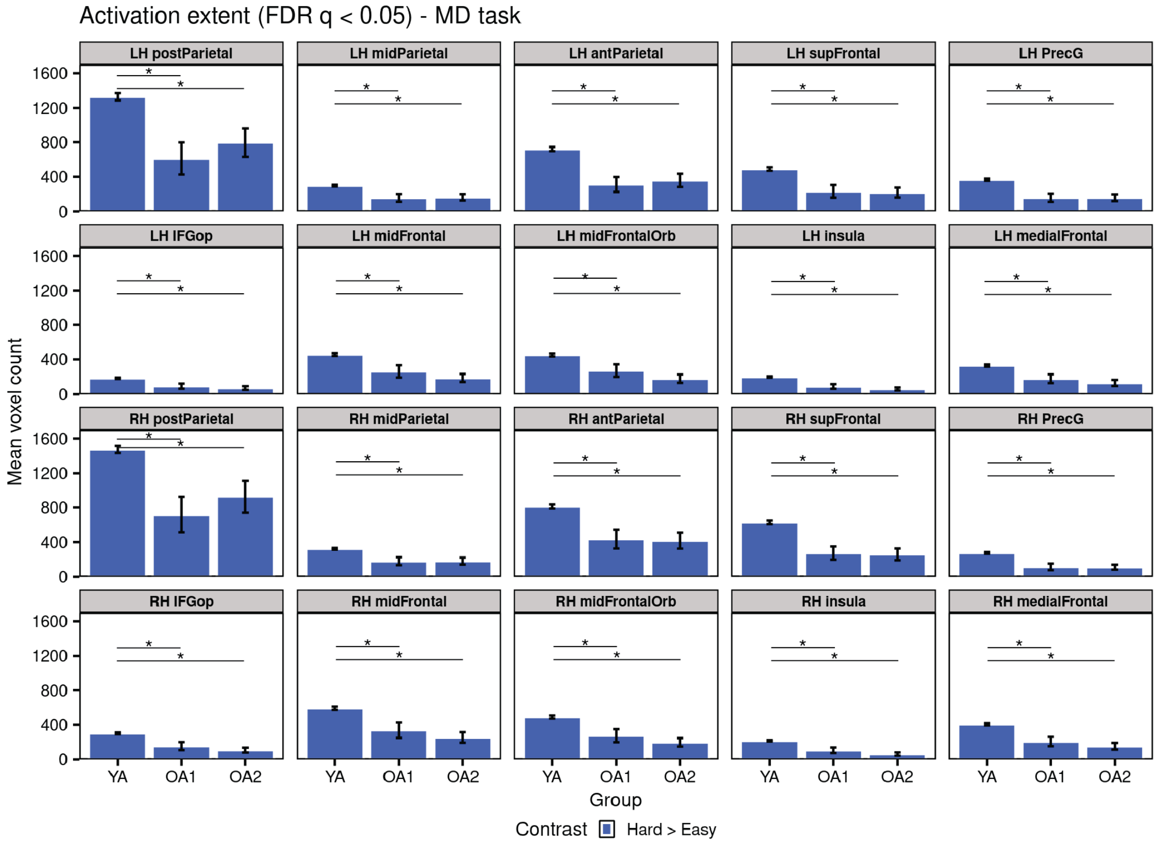


**Supp. Figure 4a. Activation extent of the language (Sentences > Nonwords lists) and the Multiple Demand (Hard > Easy) task contrasts in the younger (YA) and older adult (OA1 and OA2) cohorts within anatomical language and MD parcels.** Significant group differences are marked with an asterisk above a line (FDR-corrected *p*<0.05). LH: left-hemispheric parcel. RH: right-hemispheric parcel. IFG: inferior frontal gyrus; IFGorb: IFG pars orbitalis; MFG: middle frontal gyrus; AntTemp: anterior temporal cortex; PostTemp: posterior temporal cortex; postParietal: posterior parietal cortex; midParietal: middle parietal cortex; antParietal: anterior parietal cortex; supFrontal: superior frontal gyrus; PrecG: precentral gyrus; IFGop: IFG pars opercularis; MidFront: middle frontal gyrus; MidFrontOrb: Middle frontal gyrus, orbital part; medialFront: medial frontal cortex.

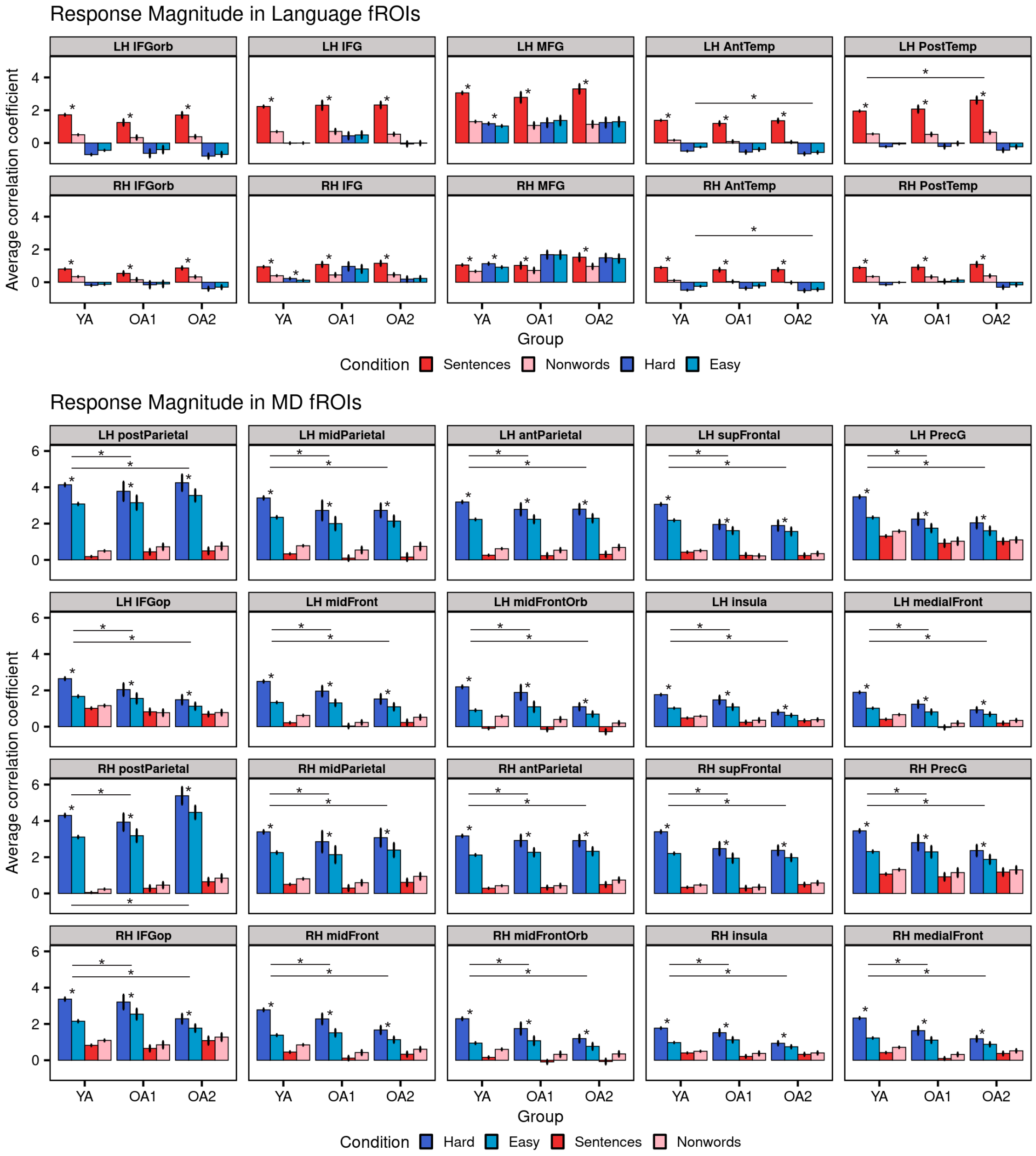


**Supp. Figure 4b. Response profiles of the language fROIs and the MD fROIs in younger (YA) and older adult (OA1 and OA2) cohorts.** Significant positive task contrasts (FDR-corrected $p<0.05$) are marked with an asterisk, significant group differences are marked with an asterisk above a line (FDR-corrected $p<0.05$). LH: left-hemispheric fROIs. RH: right-hemispheric fROIs. IFG: inferior frontal gyrus; IFGorb: IFG pars orbitalis; MFG: middle frontal gyrus; AntTemp: anterior temporal cortex; PostTemp: posterior temporal cortex; postParietal: posterior parietal cortex; midParietal: middle parietal cortex; antParietal: anterior parietal cortex; supFrontal: superior frontal gyrus; PrecG: precentral gyrus; IFGop: IFG pars opercularis; MidFront: middle frontal gyrus; MidFrontOrb: Middle frontal gyrus, orbital part; medialFront: medial frontal cortex.

### 5. Exploratory analyses of the Extended Language Network

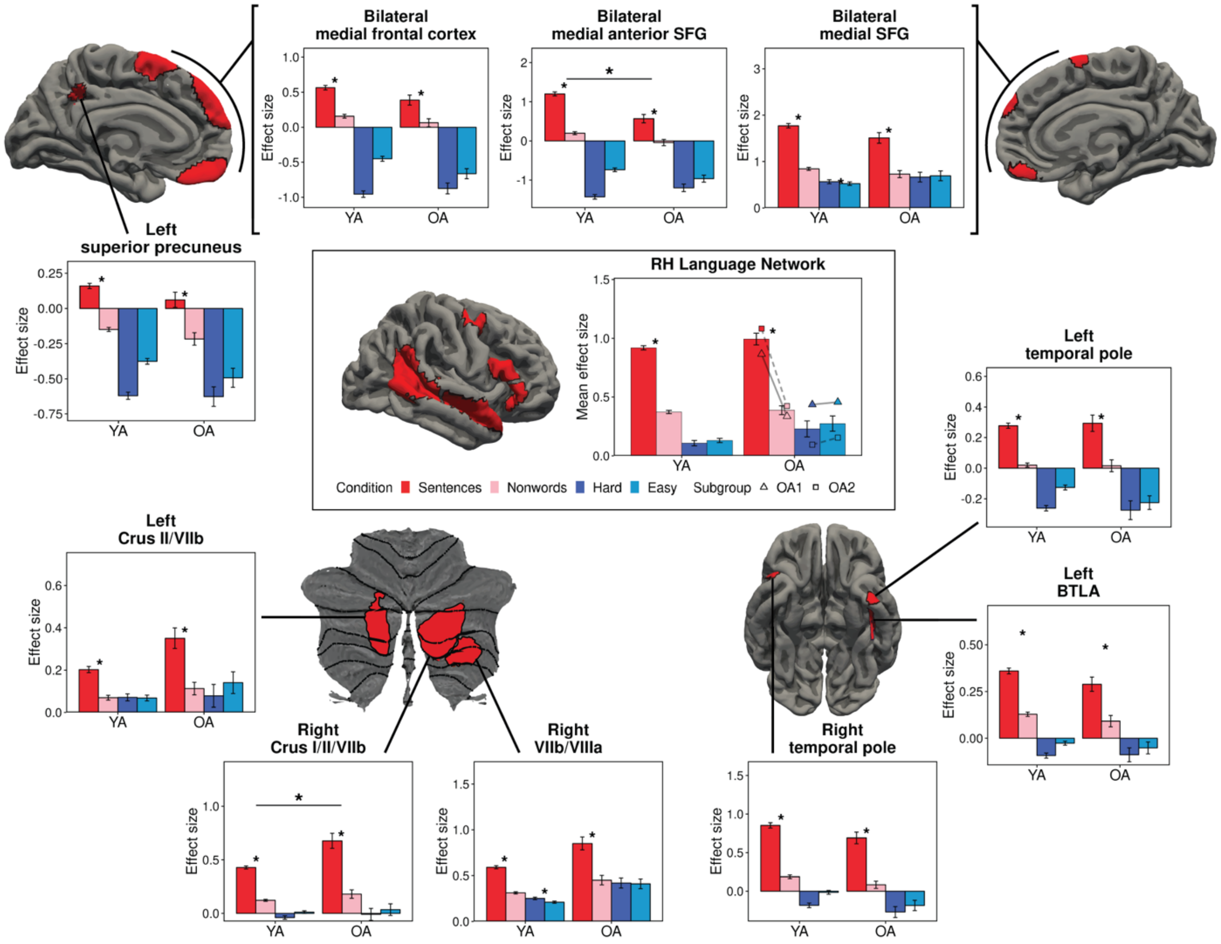


**Supp. Figure 5: Response profiles of the non-canonical language fROIs in younger and older adults**. The response to the language task contrast within the right-hemisphere (RH) language network and across most ROIs of the extended language network is similar between young adults (YA) and older adults (OA) groups, except in the bilateral medial anterior SFG where the OA group shows a lower response than the YA group, and the right Crus I/II/VIIb where the OA group shows an increased response compared to YA. The two OA subgroups in the OA group are presented in the RH plot as triangles and squares. Significant positive task contrast effects (FDR-corrected $p<0.05$) are marked with an asterisk between bars. Significant group differences in the language task contrast (FDR-corrected $p<0.05$) are marked with an asterisk above a line.

### 6. Details of the statistical models for additional results

#### Supp. Tables 6.1. Response magnitude results

##### 6.1.1. Network level: Language response in Language fROIs (LH + RH)

| Predictor | Estimate | Std. Error | Statistic | 95% CI (Lower) | 95% CI (Upper) | p-value | p.adj |
|---|---|---|---|---|---|---|---|
| (Intercept) | 0.329 | 0.014 | 24.217 | 0.303 | 0.356 | <.001 | |
| Condition = S | 0.535 | 0.006 | 85.712 | 0.523 | 0.547 | <.001 | |
| Group = OA1 | -0.048 | 0.060 | -0.807 | -0.165 | 0.069 | 0.420 | |
| Group = OA2 | -0.016 | 0.051 | -0.308 | -0.117 | 0.085 | 0.758 | |
| (Condition = S) x (Group = OA1) | 0.007 | 0.025 | 0.292 | -0.042 | 0.057 | 0.770 | 1 |
| (Condition = S) x (Group = OA2) | 0.091 | 0.027 | 3.319 | 0.037 | 0.144 | 0.001 | **0.005** |

| Predictor | Estimate | Std. Error | Statistic | 95% CI (Lower) | 95% CI (Upper) | p-value | p.adj |
|---|---|---|---|---|---|---|---|
| (Intercept) | 0.326 | 0.080 | 4.063 | 0.146 | 0.506 | 0.003 | |
| Condition = S | 0.541 | 0.008 | 71.936 | 0.527 | 0.556 | <.001 | |
| Age | -0.001 | 0.001 | -1.182 | -0.003 | 0.001 | 0.238 | |
| (Condition = S) x Age | 0.002 | 0.001 | 2.978 | 0.001 | 0.003 | 0.003 | **0.006** |

##### 6.1.2. Network level: Language response in MD fROIs (LH + RH)

| Predictor | Estimate | Std. Error | Statistic | 95% CI (Lower) | 95% CI (Upper) | p-value | p.adj |
|---|---|---|---|---|---|---|---|
| (Intercept) | 0.570 | 0.062 | 9.163 | 0.442 | 0.698 | <.001 | |
| Condition = S | -0.247 | 0.008 | -31.002 | -0.262 | -0.231 | <.001 | |
| Group = OA1 | -0.162 | 0.099 | -1.635 | -0.356 | 0.033 | 0.103 | |
| Group = OA2 | -0.046 | 0.081 | -0.562 | -0.205 | 0.114 | 0.574 | |
| (Condition = S) x (Group = OA1) | 0.053 | 0.036 | 1.468 | -0.018 | 0.123 | 0.142 | 1 |
| (Condition = S) x (Group = OA2) | 0.031 | 0.029 | 1.041 | -0.027 | 0.088 | 0.298 | 1 |

| Predictor | Estimate | Std. Error | Statistic | 95% CI (Lower) | 95% CI (Upper) | p-value | p.adj |
|---|---|---|---|---|---|---|---|
| (Intercept) | 0.559 | 0.062 | 9.061 | 0.432 | 0.687 | <.001 | |
| Condition = S | -0.242 | 0.007 | -32.369 | -0.257 | -0.228 | <.001 | |
| Age | -0.003 | 0.002 | -1.821 | -0.006 | 0.000 | 0.069 | |
| (Condition = S) x Age | 0.001 | 0.001 | 2.424 | 0.000 | 0.003 | 0.015 | 0.077 |

##### 6.1.3. Network level: MD response in Language fROIs (LH+RH)

| Predictor | Estimate | Std. Error | Statistic | 95% CI (Lower) | 95% CI (Upper) | p-value | p.adj |
|---|---|---|---|---|---|---|---|
| (Intercept) | -0.019 | 0.023 | -0.831 | -0.063 | 0.026 | 0.406 | |
| Condition = H | -0.123 | 0.006 | -19.131 | -0.135 | -0.110 | <.001 | |
| Group = OA1 | 0.172 | 0.101 | 1.692 | -0.028 | 0.371 | 0.091 | |
| Group = OA2 | -0.094 | 0.084 | -1.128 | -0.259 | 0.070 | 0.260 | |
| (Condition = H) x (Group = OA1) | 0.017 | 0.021 | 0.818 | -0.024 | 0.059 | 0.413 | 1.0 |

| | | | | | | | |
|---|---|---|---|---|---|---|---|
| (Condition = H) x (Group = OA2) | 0.018 | 0.021 | 0.864 | -0.023 | 0.059 | 0.388 | 1.0 |

| Predictor | Estimate | Std. Error | Statistic | 95% CI (Lower) | 95% CI (Upper) | p-value | p.adj |
|---|---|---|---|---|---|---|---|
| (Intercept) | -0.018 | 0.164 | -0.108 | -0.387 | 0.352 | 0.917 | |
| Condition = H | -0.121 | 0.013 | -9.146 | -0.147 | -0.095 | <.001 | |
| Age | 0.001 | 0.002 | 0.518 | -0.003 | 0.004 | 0.605 | |
| (Condition = H) x Age | 0.001 | 0.001 | 1.114 | -0.001 | 0.003 | 0.265 | 1 |

**6.1.4. Hemisphere level: MD response in the MD LH fROIs**

| Predictor | Estimate | Std. Error | Statistic | 95% CI (Lower) | 95% CI (Upper) | p-value | p.adj |
|---|---|---|---|---|---|---|---|
| (Intercept) | 1.264 | 0.025 | 49.637 | 1.214 | 1.314 | <.001 | |
| Condition = H | 0.551 | 0.006 | 90.805 | 0.539 | 0.563 | <.001 | |
| Group = OA1 | -0.100 | 0.122 | -0.820 | -0.338 | 0.139 | 0.412 | |
| Group = OA2 | -0.221 | 0.094 | -2.354 | -0.405 | -0.037 | 0.019 | |
| (Condition = H) x (Group = OA1) | -0.264 | 0.027 | -9.899 | -0.316 | -0.212 | <.001 | **<.001** |
| (Condition = H) x (Group = OA2) | -0.337 | 0.020 | -17.021 | -0.376 | -0.298 | <.001 | **<.001** |

| Predictor | Estimate | Std. Error | Statistic | 95% CI (Lower) | 95% CI (Upper) | p-value | p.adj |
|---|---|---|---|---|---|---|---|
| (Intercept) | 1.244 | 0.135 | 9.248 | 0.943 | 1.546 | <.001 | |
| Condition = H | 0.516 | 0.010 | 50.159 | 0.496 | 0.536 | <.001 | |
| Age | -0.004 | 0.002 | -1.977 | -0.008 | 0.000 | 0.048 | |
| (Condition = H) x Age | -0.009 | 0.001 | -10.738 | -0.010 | -0.007 | <.001 | **<.001** |

**6.1.5. Hemisphere level: MD response in the MD RH fROIs**

| Predictor | Estimate | Std. Error | Statistic | 95% CI (Lower) | 95% CI (Upper) | p-value | p.adj |
|---|---|---|---|---|---|---|---|
| (Intercept) | 1.293 | 0.028 | 46.934 | 1.239 | 1.347 | <.001 | |
| Condition = H | 0.623 | 0.006 | 97.510 | 0.610 | 0.635 | <.001 | |
| Group = OA1 | 0.025 | 0.131 | 0.192 | -0.233 | 0.283 | 0.848 | |
| Group = OA2 | -0.078 | 0.102 | -0.766 | -0.278 | 0.122 | 0.444 | |
| (Condition = H) x (Group = OA1) | -0.311 | 0.027 | -11.638 | -0.364 | -0.259 | <.001 | **<.001** |
| (Condition = H) x (Group = OA2) | -0.359 | 0.023 | -15.408 | -0.404 | -0.313 | <.001 | **<.001** |

| Predictor | Estimate | Std. Error | Statistic | 95% CI (Lower) | 95% CI (Upper) | p-value | p.adj |
|---|---|---|---|---|---|---|---|
| (Intercept) | 1.289 | 0.131 | 9.807 | 0.995 | 1.582 | <.001 | |
| Condition = H | 0.584 | 0.011 | 54.210 | 0.563 | 0.605 | <.001 | |
| Age | -0.001 | 0.002 | -0.270 | -0.005 | 0.004 | 0.787 | |
| (Condition = H) x Age | -0.010 | 0.001 | -11.353 | -0.011 | -0.008 | <.001 | **<.001** |

### Supp. Tables 6.2. Activation extent results.

#### 6.2.1. Extent of Whole-Brain Activation Maps (FDR *p*<0.05)

##### 6.2.1.1. Language Contrast S>N

| Predictor | Estimate | Std. Error | Statistic | 95% CI (Lower) | 95% CI (Upper) | p-value | p.adj |
|---|---|---|---|---|---|---|---|
| (Intercept) | 109.936 | 1.586 | 69.311 | 106.820 | 113.051 | <.001 | |
| Group = OA1 | 2.465 | 7.018 | 0.351 | -11.320 | 16.251 | 0.726 | 0.726 |
| Group = OA2 | 9.374 | 5.873 | 1.596 | -2.162 | 20.911 | 0.111 | 0.333 |

| Predictor | Estimate | Std. Error | Statistic | 95% CI (Lower) | 95% CI (Upper) | p-value | p.adj |
|---|---|---|---|---|---|---|---|
| (Intercept) | 110.702 | 1.491 | 74.254 | 107.773 | 113.630 | <.001 | |
| Age | 0.133 | 0.116 | 1.147 | -0.095 | 0.362 | 0.252 | 0.252 |

##### 6.2.1.2. MD Contrast: H>E

| Predictor | Estimate | Std. Error | Statistic | 95% CI (Lower) | 95% CI (Upper) | p-value | p.adj |
|---|---|---|---|---|---|---|---|
| (Intercept) | 154.742 | 2.085 | 74.205 | 150.646 | 158.838 | <.001 | |
| Group = OA1 | -11.829 | 11.929 | -0.992 | -35.262 | 11.603 | 0.322 | 0.644 |
| Group = OA2 | -22.178 | 9.715 | -2.283 | -41.261 | -3.095 | 0.023 | 0.091 |

| Predictor | Estimate | Std. Error | Statistic | 95% CI (Lower) | 95% CI (Upper) | p-value | p.adj |
|---|---|---|---|---|---|---|---|
| (Intercept) | 152.641 | 2.027 | 75.310 | 148.659 | 156.622 | <.001 | |
| Age | -0.495 | 0.158 | -3.132 | -0.806 | -0.185 | 0.002 | **0.004** |

#### 6.2.2. Extent of Activation Maps within Parcels (thresholded at FDR-corrected *p*<0.05)

##### 6.2.2.1. Language Contrast: S>N (LH+RH)

| Predictor | Estimate | Std. Error | Statistic | 95% CI (Lower) | 95% CI (Upper) | p-value | p.adj |
|---|---|---|---|---|---|---|---|
| (Intercept) | 12.265 | 2.499 | 4.908 | 6.626 | 17.904 | <.001 | |
| Group = OA1 | 0.225 | 1.053 | 0.214 | -1.843 | 2.293 | 0.831 | 1.000 |
| Group = OA2 | 1.439 | 0.881 | 1.634 | -0.291 | 3.170 | 0.103 | 0.607 |

| Predictor | Estimate | Std. Error | Statistic | 95% CI (Lower) | 95% CI (Upper) | p-value | p.adj |
|---|---|---|---|---|---|---|---|
| (Intercept) | 12.375 | 2.498 | 4.954 | 6.737 | 18.013 | <.001 | |
| Age | 0.027 | 0.017 | 1.526 | -0.008 | 0.061 | 0.128 | 0.315 |

##### 6.2.2.2. MD Contrast: H>E (LH)

| Predictor | Estimate | Std. Error | Statistic | 95% CI (Lower) | 95% CI (Upper) | p-value | p.adj |
|---|---|---|---|---|---|---|---|
| (Intercept) | 18.370 | 1.935 | 9.496 | 14.057 | 22.683 | <.001 | |
| Group = OA1 | -8.230 | 1.953 | -4.213 | -12.067 | -4.393 | <.001 | **<.001** |
| Group = OA2 | -9.008 | 1.605 | -5.614 | -12.160 | -5.857 | <.001 | **<.001** |

| Predictor | Estimate | Std. Error | Statistic | 95% CI (Lower) | 95% CI (Upper) | p-value | p.adj |
|---|---|---|---|---|---|---|---|
| (Intercept) | 17.360 | 1.929 | 9.001 | 13.053 | 21.668 | <.001 | |
| Age | -0.220 | 0.032 | -6.921 | -0.282 | -0.157 | <.001 | **<.001** |

##### 6.2.2.3. MD Contrast: H>E (RH)

| Predictor | Estimate | Std. Error | Statistic | 95% CI (Lower) | 95% CI (Upper) | p-value | p.adj |
|---|---|---|---|---|---|---|---|
| (Intercept) | 20.066 | 2.069 | 9.700 | 15.447 | 24.686 | <.001 | |

| Predictor | Estimate | Std. Error | Statistic | 95% CI (Lower) | 95% CI (Upper) | p-value | p.adj |
|---|---|---|---|---|---|---|---|
| Group = OA1 | -8.901 | 1.971 | -4.516 | -12.773 | -5.030 | <.001 | **<.001** |
| Group = OA2 | -9.879 | 1.619 | -6.102 | -13.060 | -6.699 | <.001 | **<.001** |

| Predictor | Estimate | Std. Error | Statistic | 95% CI (Lower) | 95% CI (Upper) | p-value | p.adj |
|---|---|---|---|---|---|---|---|
| (Intercept) | 18.965 | 2.063 | 9.192 | 14.351 | 23.579 | <.001 | |
| Age | -0.242 | 0.032 | -7.568 | -0.305 | -0.179 | <.001 | **<.001** |

### Supp. Tables 6.3. Functional connectivity results

#### 6.3.1. Resting-State: Within Language Network

##### 6.3.1.1. Network Level (All Connections)

| Predictor | Estimate | Std. Error | Statistic | 95% CI (Lower) | 95% CI (Upper) | p-value | p.adj |
|---|---|---|---|---|---|---|---|
| (Intercept) | 0.334 | 0.031 | 10.897 | 0.272 | 0.396 | <.001 | |
| Group = OA1 | 0.015 | 0.034 | 0.432 | -0.052 | 0.082 | 0.666 | 1.000 |
| Group = OA2 | 0.058 | 0.025 | 2.350 | 0.009 | 0.107 | 0.020 | 0.142 |

| Predictor | Estimate | Std. Error | Statistic | 95% CI (Lower) | 95% CI (Upper) | p-value | p.adj |
|---|---|---|---|---|---|---|---|
| (Intercept) | 0.352 | 0.029 | 11.969 | 0.292 | 0.411 | <.001 | |
| Age | 0.001 | 0.001 | 1.545 | 0.000 | 0.002 | 0.125 | 0.374 |

##### 6.3.1.2. Hemisphere Level (Inter-hemispheric)

| Predictor | Estimate | Std. Error | Statistic | 95% CI (Lower) | 95% CI (Upper) | p-value | p.adj |
|---|---|---|---|---|---|---|---|
| (Intercept) | 0.270 | 0.029 | 9.406 | 0.212 | 0.328 | <.001 | |
| Group = OA1 | 0.031 | 0.037 | 0.863 | -0.041 | 0.104 | 0.390 | 1.000 |
| Group = OA2 | 0.076 | 0.027 | 2.851 | 0.023 | 0.129 | 0.005 | **0.040** |

| Predictor | Estimate | Std. Error | Statistic | 95% CI (Lower) | 95% CI (Upper) | p-value | p.adj |
|---|---|---|---|---|---|---|---|
| (Intercept) | 0.295 | 0.027 | 10.865 | 0.240 | 0.351 | <.001 | |
| Age | 0.001 | 0.001 | 2.014 | 0.000 | 0.003 | 0.046 | 0.184 |

#### 6.3.2. Story Listening: Within Language Network

##### 6.3.2.1. Network Level (All Connections)

| Predictor | Estimate | Std. Error | Statistic | 95% CI (Lower) | 95% CI (Upper) | p-value | p.adj |
|---|---|---|---|---|---|---|---|
| (Intercept) | 0.501 | 0.043 | 11.764 | 0.414 | 0.589 | <.001 | |
| Group = OA2 | 0.035 | 0.034 | 1.036 | -0.032 | 0.103 | 0.302 | 0.907 |

| Predictor | Estimate | Std. Error | Statistic | 95% CI (Lower) | 95% CI (Upper) | p-value | p.adj |
|---|---|---|---|---|---|---|---|
| (Intercept) | 0.513 | 0.041 | 12.423 | 0.428 | 0.597 | <.001 | |
| Age | 0.000 | 0.001 | 0.555 | -0.001 | 0.002 | 0.580 | 1.000 |

##### 6.3.2.2. Hemisphere Level (Inter-hemispheric)

| Predictor | Estimate | Std. Error | Statistic | 95% CI (Lower) | 95% CI (Upper) | p-value | p.adj |
|---|---|---|---|---|---|---|---|
| (Intercept) | 0.434 | 0.040 | 10.872 | 0.353 | 0.515 | <.001 | |

| Predictor | Estimate | Std. Error | Statistic | 95% CI (Lower) | 95% CI (Upper) | p-value | p.adj |
|---|---|---|---|---|---|---|---|
| Group = OA2 | 0.065 | 0.036 | 1.812 | -0.006 | 0.136 | 0.073 | 0.290 |

| Predictor | Estimate | Std. Error | Statistic | 95% CI (Lower) | 95% CI (Upper) | p-value | p.adj |
|---|---|---|---|---|---|---|---|
| (Intercept) | 0.454 | 0.038 | 11.860 | 0.376 | 0.533 | <.001 | |
| Age | 0.001 | 0.001 | 1.293 | -0.001 | 0.003 | 0.199 | 0.795 |

#### 6.3.3. Resting-State: Within MD Network

##### 6.3.3.1. Network Level (All Connections)

| Predictor | Estimate | Std. Error | Statistic | 95% CI (Lower) | 95% CI (Upper) | p-value | p.adj |
|---|---|---|---|---|---|---|---|
| (Intercept) | 0.320 | 0.018 | 18.142 | 0.285 | 0.355 | <.001 | |
| Group = OA1 | -0.071 | 0.030 | -2.397 | -0.130 | -0.012 | 0.018 | 0.054 |
| Group = OA2 | -0.090 | 0.022 | -4.146 | -0.133 | -0.047 | <.001 | **<.001** |

| Predictor | Estimate | Std. Error | Statistic | 95% CI (Lower) | 95% CI (Upper) | p-value | p.adj |
|---|---|---|---|---|---|---|---|
| (Intercept) | 0.287 | 0.016 | 18.189 | 0.256 | 0.318 | <.001 | |
| Age | -0.003 | 0.000 | -5.202 | -0.004 | -0.002 | <.001 | **<.001** |

##### 6.3.3.2. Hemisphere Level (Inter-hemispheric)

| Predictor | Estimate | Std. Error | Statistic | 95% CI (Lower) | 95% CI (Upper) | p-value | p.adj |
|---|---|---|---|---|---|---|---|
| (Intercept) | 0.250 | 0.016 | 15.334 | 0.218 | 0.282 | <.001 | |
| Group = OA1 | -0.045 | 0.030 | -1.481 | -0.105 | 0.015 | 0.141 | 0.141 |
| Group = OA2 | -0.065 | 0.022 | -2.939 | -0.109 | -0.021 | 0.004 | **0.017** |

| Predictor | Estimate | Std. Error | Statistic | 95% CI (Lower) | 95% CI (Upper) | p-value | p.adj |
|---|---|---|---|---|---|---|---|
| (Intercept) | 0.226 | 0.014 | 15.947 | 0.198 | 0.254 | <.001 | |
| Age | -0.002 | 0.001 | -3.651 | -0.003 | -0.001 | <.001 | **<.001** |

#### 6.3.4. Story Listening: Within MD Network

##### 6.3.4.1. Network Level (All Connections)

| Predictor | Estimate | Std. Error | Statistic | 95% CI (Lower) | 95% CI (Upper) | p-value | p.adj |
|---|---|---|---|---|---|---|---|
| (Intercept) | 0.301 | 0.016 | 19.049 | 0.270 | 0.332 | <.001 | |
| Group = OA2 | -0.063 | 0.019 | -3.352 | -0.101 | -0.026 | 0.001 | **0.002** |

| Predictor | Estimate | Std. Error | Statistic | 95% CI (Lower) | 95% CI (Upper) | p-value | p.adj |
|---|---|---|---|---|---|---|---|
| (Intercept) | 0.281 | 0.015 | 19.330 | 0.252 | 0.310 | <.001 | |
| Age | -0.002 | 0.000 | -3.751 | -0.003 | -0.001 | <.001 | **<.001** |

##### 6.3.4.2. Hemisphere Level (Inter-hemispheric)

| Predictor | Estimate | Std. Error | Statistic | 95% CI (Lower) | 95% CI (Upper) | p-value | p.adj |
|---|---|---|---|---|---|---|---|
| (Intercept) | 0.230 | 0.015 | 15.838 | 0.202 | 0.259 | <.001 | |
| Group = OA2 | -0.030 | 0.020 | -1.500 | -0.069 | 0.010 | 0.136 | 0.136 |

| Predictor | Estimate | Std. Error | Statistic | 95% CI (Lower) | 95% CI (Upper) | p-value | p.adj |
|---|---|---|---|---|---|---|---|
| (Intercept) | 0.221 | 0.013 | 16.978 | 0.195 | 0.246 | <.001 | |
| Age | -0.001 | 0.000 | -1.929 | -0.002 | 0.000 | 0.056 | 0.056 |

#### 6.3.5. Resting-State: Between Language and MD Networks

##### 6.3.5.1. Network Level (All Connections)

| Predictor | Estimate | Std. Error | Statistic | 95% CI (Lower) | 95% CI (Upper) | p-value | p.adj |
|---|---|---|---|---|---|---|---|
| (Intercept) | 0.018 | 0.015 | 1.238 | -0.011 | 0.047 | 0.218 | |
| Group = OA1 | 0.020 | 0.022 | 0.892 | -0.024 | 0.064 | 0.374 | 1.000 |
| Group = OA2 | 0.081 | 0.016 | 4.974 | 0.048 | 0.113 | <.001 | **<.001** |

| Predictor | Estimate | Std. Error | Statistic | 95% CI (Lower) | 95% CI (Upper) | p-value | p.adj |
|---|---|---|---|---|---|---|---|
| (Intercept) | 0.043 | 0.014 | 3.106 | 0.015 | 0.070 | 0.002 | |
| Age | 0.002 | 0.000 | 3.808 | 0.001 | 0.002 | <.001 | **<.001** |

##### 6.3.5.2. Hemisphere Level (Inter-hemispheric)

| Predictor | Estimate | Std. Error | Statistic | 95% CI (Lower) | 95% CI (Upper) | p-value | p.adj |
|---|---|---|---|---|---|---|---|
| (Intercept) | -0.031 | 0.013 | -2.483 | -0.056 | -0.006 | 0.014 | |
| Group = OA1 | 0.017 | 0.023 | 0.769 | -0.027 | 0.062 | 0.443 | 1.000 |
| Group = OA2 | 0.082 | 0.016 | 4.984 | 0.049 | 0.114 | <.001 | **<.001** |

| Predictor | Estimate | Std. Error | Statistic | 95% CI (Lower) | 95% CI (Upper) | p-value | p.adj |
|---|---|---|---|---|---|---|---|
| (Intercept) | -0.007 | 0.011 | -0.607 | -0.030 | 0.016 | 0.545 | |
| Age | 0.002 | 0.000 | 3.762 | 0.001 | 0.002 | <.001 | **<.001** |

#### 6.3.6. Story Listening: Between Language and MD Networks

##### 6.3.6.1. Network Level (All Connections)

| Predictor | Estimate | Std. Error | Statistic | 95% CI (Lower) | 95% CI (Upper) | p-value | p.adj |
|---|---|---|---|---|---|---|---|
| (Intercept) | 0.013 | 0.017 | 0.764 | -0.020 | 0.046 | 0.447 | |
| Group = OA2 | 0.034 | 0.018 | 1.937 | -0.001 | 0.070 | 0.055 | 0.166 |

| Predictor | Estimate | Std. Error | Statistic | 95% CI (Lower) | 95% CI (Upper) | p-value | p.adj |
|---|---|---|---|---|---|---|---|
| (Intercept) | 0.024 | 0.016 | 1.504 | -0.008 | 0.055 | 0.136 | |
| Age | 0.001 | 0.000 | 1.789 | 0.000 | 0.002 | 0.076 | 0.229 |

##### 6.3.6.2. Hemisphere Level (Inter-hemispheric)

| Predictor | Estimate | Std. Error | Statistic | 95% CI (Lower) | 95% CI (Upper) | p-value | p.adj |
|---|---|---|---|---|---|---|---|
| (Intercept) | -0.027 | 0.015 | -1.872 | -0.056 | 0.002 | 0.063 | |
| Group = OA2 | 0.044 | 0.018 | 2.464 | 0.009 | 0.079 | 0.015 | 0.061 |

| Predictor | Estimate | Std. Error | Statistic | 95% CI (Lower) | 95% CI (Upper) | p-value | p.adj |
|---|---|---|---|---|---|---|---|
| (Intercept) | -0.014 | 0.014 | -1.002 | -0.040 | 0.013 | 0.319 | |
| Age | 0.001 | 0.000 | 2.160 | 0.000 | 0.002 | 0.033 | 0.131 |

#### Supp. Tables 6.4. Spatial activation overlap results within domain between runs.

**6.4.1. Within Language Network**

| Predictor | Estimate | Std. Error | Statistic | 95% CI (Lower) | 95% CI (Upper) | p-value | p.adj |
|---|---|---|---|---|---|---|---|
| (Intercept) | -0.804 | 0.033 | -24.254 | -0.869 | -0.739 | <.001 | |
| Group = OA1 | -0.143 | 0.141 | -1.016 | -0.420 | 0.133 | 0.310 | 0.819 |
| Group = OA2 | -0.006 | 0.120 | -0.046 | -0.241 | 0.230 | 0.963 | 0.963 |

| Predictor | Estimate | Std. Error | Statistic | 95% CI (Lower) | 95% CI (Upper) | p-value | p.adj |
|---|---|---|---|---|---|---|---|
| (Intercept) | -0.811 | 0.031 | -25.976 | -0.873 | -0.750 | <.001 | |
| Age | 0.000 | 0.002 | -0.122 | -0.005 | 0.004 | 0.903 | 0.991 |

**6.4.2 Within MD Network**

| Predictor | Estimate | Std. Error | Statistic | 95% CI (Lower) | 95% CI (Upper) | p-value | p.adj |
|---|---|---|---|---|---|---|---|
| (Intercept) | -0.736 | 0.043 | -17.136 | -0.820 | -0.651 | <.001 | |
| Group = OA1 | -0.281 | 0.181 | -1.557 | -0.636 | 0.073 | 0.119 | 0.597 |
| Group = OA2 | -0.696 | 0.162 | -4.293 | -1.014 | -0.378 | <.001 | **<.001** |

| Predictor | Estimate | Std. Error | Statistic | 95% CI (Lower) | 95% CI (Upper) | p-value | p.adj |
|---|---|---|---|---|---|---|---|
| (Intercept) | -0.799 | 0.041 | -19.552 | -0.879 | -0.719 | <.001 | |
| Age | -0.015 | 0.003 | -4.807 | -0.021 | -0.009 | <.001 | **<.001** |
